# A novel TGF-β-responsive enhancer regulates SRC expression and epithelial–mesenchymal transition-associated cell migration

**DOI:** 10.1101/2022.11.25.517908

**Authors:** Soshi Noshita, Yuki Kubo, Kentaro Kajiwara, Daisuke Okuzaki, Shigeyuki Nada, Masato Okada

**Author notes:** To whom correspondence should be addressed: Masato Okada.

## Abstract

The non-receptor tyrosine kinase SRC is overexpressed and/or hyperactivated in various human cancers, and facilitates cancer progression by promoting invasion and metastasis. However, the mechanisms underlying SRC upregulation are poorly understood. In this study, we demonstrated that transforming growth factor-β (TGF-β) induces SRC expression at the transcriptional level by activating a novel intragenic SRC enhancer. In the human breast epithelial cell line MCF10A, TGF-β1 stimulation upregulated the *SRC1A* promoter, resulting in increased SRC mRNA and protein levels. Chromatin immunoprecipitation (ChIP)-sequencing analysis revealed that SMAD complex is recruited to three enhancer regions ~15 kb upstream and downstream of the *SRC* promoter, and one of them is capable of activating the *SRC* promoter in response to TGF-β. In addition, JUN, which is a member of the activator protein (AP)-1 family, also localizes to the enhancer and regulates TGF-β-induced SRC expression. Furthermore, the total amount of active SRC also increased, coinciding with the TGF-β-induced SRC expression. In TGF-β-induced epithelial–mesenchymal transition (EMT), TGF-β-induced SRC upregulation plays a crucial role in EMT-associated cell migration by activating the SRC/focal adhesion kinase (FAK) circuit. Overall, these results suggest that TGF-β-induced SRC upregulation promotes cancer cell invasion and metastasis in a subset of human malignancies.

## Introduction

The proto-oncogene product SRC, a non-receptor tyrosine kinase, plays a pivotal role in the regulation of various cellular functions, including proliferation, survival, adhesion, and migration. SRC facilitates invasion and metastasis in cancer cells by accelerating these functions (Guarino, 2010; Yeatman, 2004). SRC is overexpressed and/or activated in various human malignancies, implying that SRC plays a crucial role in tumor progression (Ishizawar and Parsons, 2004; RB and TJ, 2000; Verbeek et al., 1996; Wiener et al., 2003; Yeatman, 2004). Overexpression of SRC was observed over two decades ago in human breast cancer (Verbeek et al., 1996). Furthermore, a current multi-omics study also revealed that SRC is overexpressed at the mRNA level and is an essential gene for breast cancer progression (López-Cortés et al., 2020). However, the mechanisms underlying SRC overexpression in breast cancer still remain unclear. In particular, transcriptional regulation of *SRC* is poorly understood.

Transforming growth factor-β (TGF-β), a multifunctional cytokine, regulates multiple cellular functions, including cell proliferation, differentiation, and motility, to maintain tissue homeostasis (Massagué, 2012). Once TGF-β binds to the TGF-β type II receptor, the activated TGF-β type II receptor phosphorylates the TGF-β type I receptor to activate downstream signaling, such as phosphorylation of SMAD2/3. Phosphorylated SMAD2/3 forms a heterotrimeric complex with SMAD4 (Shi and Massagué, 2003). The SMAD2/3-SMAD4 complex is transported into the nucleus and regulates the expression of target genes by directly binding to regulatory gene sequences in cooperation with other transcription factors and/or co-activators/repressors (SMAD pathway). TGF-β also activates other signaling pathways, including the extracellular signal-regulated kinase (ERK), c-Jun N-terminal kinase (JNK), and phosphoinositide 3-kinase (PI3K) pathways (non-SMAD pathways) (Colak and ten Dijke, 2017; Gaarenstroom and Hill, 2014).

TGF-β plays a dual role in cancer as a tumor suppressor and promoter. As a tumor suppressor, TGF-β induces cell cycle arrest and apoptosis in normal and premalignant cells to maintain tissue homeostasis. In cancer cells, however, TGF-β promotes tumor progression, including invasion and metastasis, by inducing epithelial–mesenchymal transition (EMT) (Colak and ten Dijke, 2017; Padua and Massagué, 2009). During EMT, epithelial cells lose cell–cell adhesion and apical-basal polarity and acquire mesenchymal traits such as cell motility and invasiveness (Fuxe et al., 2020). In addition to TGF-β, SRC promotes cell migration and invasion by regulating the turnover of focal adhesion, cytoskeletal remodeling, formation of invadopodia, and secretion of matrix metalloproteinases (MMPs) to degrade the extracellular matrix (ECM). Therefore, further studies on the relationship between TGF-β signaling and SRC should be conducted to gain a deeper understanding of cancer progression.

In previous studies, we found that ARHGEF5, a Rho guanine nucleotide exchange factor, is upregulated and promotes cell migration by activating the Rho-ROCK pathway during TGF-β-induced EMT in the human breast epithelial cell line MCF10A (Komiya et al., 2016; Kuroiwa et al., 2011). Moreover, SRC facilitates the activation of ARHGEF5, leading to cell invasion and tumor growth (Komiya et al., 2016). In addition to these findings, we found that SRC protein and mRNA levels increased, coinciding with TGF-β-induced EMT. However, the mechanisms by which TGF-β signaling regulates SRC gene expression remains unknown.

In this study, we found that a TGF-β-responsive enhancer located in the *SRC* intragenic region upregulates SRC promoter activity via the TGF-β/SMAD and TGF-β/JNK/JUN pathways. We also demonstrated that increased SRC expression is essential for promoting EMT-associated cell motility by phosphorylating FAK. This is the first report to demonstrate the transcriptional regulation of *SRC* induced by extracellular stimulation, and these findings help elucidate how SRC is overexpressed in various types of human cancer.

## Results

### SRC protein and exon1A transcripts are increased during TGF-β-induced EMT

To analyze SRC expression during EMT, we used the normal human breast epithelial cell line, MCF10A. TGF-β stimulation of MCF10A cells induced morphological changes, such as the reduction of E-cadherin-mediated cell–cell adhesion and the formation of stress fibers (Fig. 1A). In addition, switching from E-cadherin to N-cadherin was observed, indicating that EMT was induced in these cells (Fig. 1B). During EMT, SRC protein and mRNA levels were elevated by 2- to 4-fold in a time-dependent manner (Fig. 1B and C). In addition to MCF10A cells, TGF-β-induced SRC expression was also observed in the triple-negative breast cancer cell line BT-549 (Fig. S1A and B). SRC has been reported to be overexpressed in renal cancer cell lines by downregulating miR-205, which targets and degrades SRC mRNA (Majid et al., 2011). To verify whether TGF-β affects the stability of SRC mRNA, we determined the degradation rate of SRC mRNA in the presence or absence of TGF-β by arresting transcription using actinomycin D. The result showed that TGF-β treatment did not significantly affect the stability of *SRC* mRNA (Fig. 1D), suggesting that TGF-β regulates *SRC* expression at the transcriptional level.

**Figure 1.**
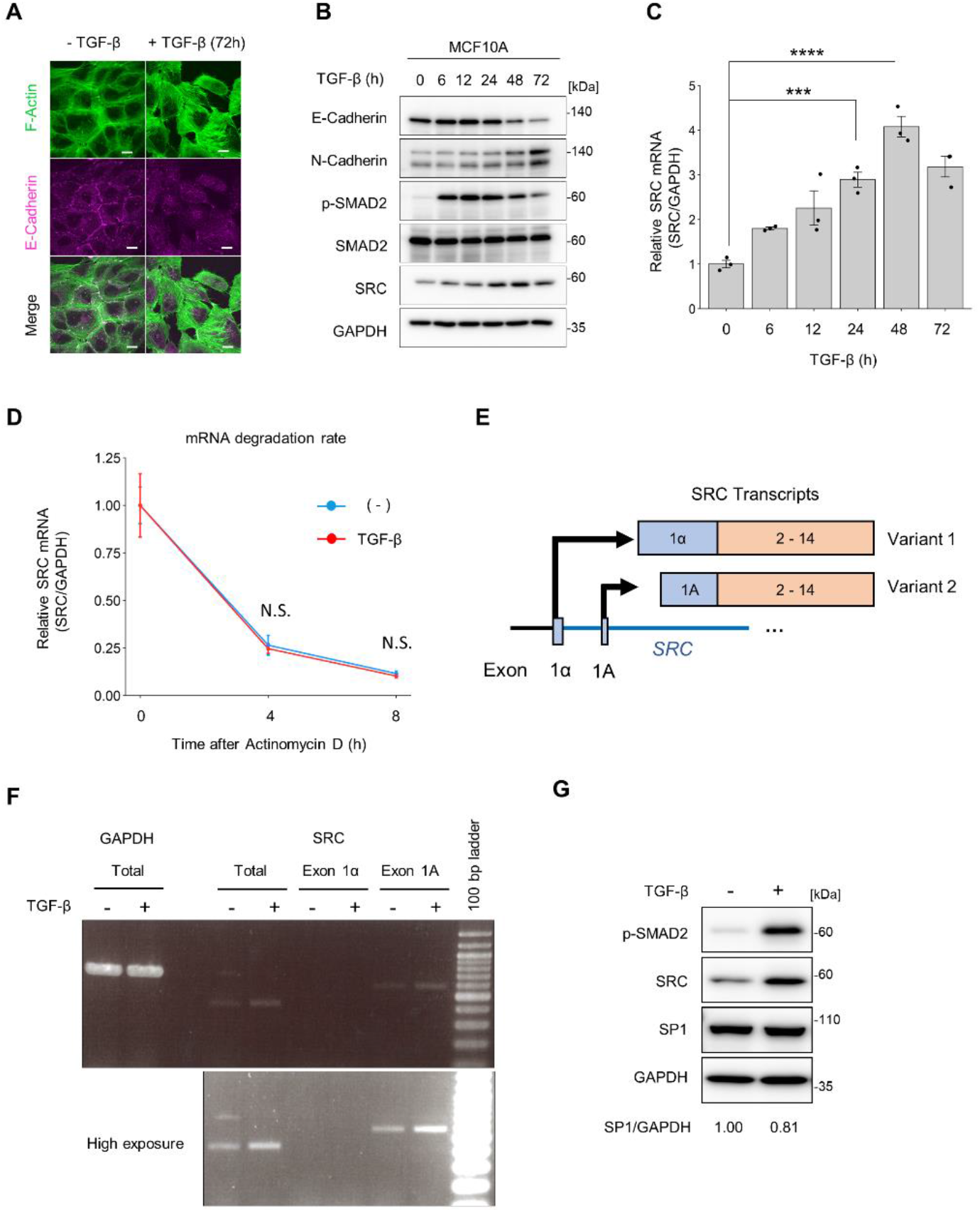
TGF-β stimulation upregulates *SRC* transcription in MCF10A. (A) MCF10A cells were treated with TGF-β1 (10 ng/ml) for 72h. The cells were subjected to immunofluorescence staining for F-actin and E-cadherin. The scale bar represents 10 μm. (B, C) MCF10A cells were treated with TGF-β1(10 ng/ml) for the indicated times. (B) Cell lysates were subjected to immunoblotting using the indicated antibodies. (C) Total RNA was isolated and subjected to qPCR. Relative expression levels were calculated using the mean value at 0h. (D) MCF10A cells were treated with actinomycin D (10 μg/ml) for 4 or 8h in the presence or absence of TGF-β1(10 ng/ml). TGF-β stimulation was conducted for 24h, and then subjected to quantitative real-time PCR. Relative expression levels were calculated using the mean value at 0h. (E) Schematic diagram of the transcriptional variants of SRC mRNA. Each variant is regulated by two different promoters and contains different exon1. (F) Transcriptional variants of SRC mRNA were analyzed by real-time PCR. (G) Immunoblotting analysis for the expression of SP1 in MCF10A cells. (F and G) MCF10A cells were treated with TGF-β1(10 ng/ml) for 24h. (C and D) The mean ratios ± SEs were obtained from three independent experiments. ***, *p* < 0.001; ****, *p* < 0.0001; N.S., not significantly different; One-way ANOVA with Tukey’s post hoc test.

There are two types of *SRC* transcripts whose expression is regulated by two different promoters: SRC1A and SRC1α (Bonham et al., 2000; Ritchie et al., 2000) (Fig. 1E). Reverse transcription PCR (RT-PCR) analysis showed that SRC exon1A transcripts, but not SRC exon1α transcripts, were increased by TGF-β stimulation, implying that the SRC1A promoter was selectively activated (Fig. 1F). Previous studies revealed that SRC1A promoter activity is dependent on the SP1 transcription factor, and the increase in SP1 results in SRC1A promoter activation and SRC expression (Liu et al., 2018; Ritchie et al., 2000). However, TGF-β stimulation did not induce SP1 expression in MCF10A cells (Fig. 1G). Taken together, these results suggested that there are unknown regulatory mechanisms underlying TGF-β-induced SRC transcription.

### SRC expression is directly regulated via TGF-β/SMAD canonical signaling pathway

To identify the signaling pathways and transcription factors that induce *SRC* promoter activation, we first examined the TGF-β/SMAD signaling pathway. The TGFβRI inhibitor LY364947 inhibited C-terminal phosphorylation of SMAD2 and TGF-β-induced SRC protein expression (Fig. 2A). We also established SMAD4-knockout (KO) MCF10A cells using CRISPR/Cas9. TGF-β failed to induce SRC expression at the protein and mRNA levels in SMAD4-KO cells, indicating that the TGF-β/SMAD pathway is essential for the regulation of SRC transcription (Fig. 2B and C). As TGF-β drives the EMT process by inducing various EMT-associated transcription factors (Fuxe et al., 2020), we also examined the contribution of TGF-β-induced factors. Although inhibition of *de novo* protein synthesis using cycloheximide reduced basal levels of SRC mRNA, TGF-β-induced elevation of SRC transcription was observed even in the presence of cycloheximide and was suppressed by LY364947(Fig. 2D). These results suggest that SRC transcription is regulated by the preexisting TGF-β/SMAD pathway.

**Figure 2.**
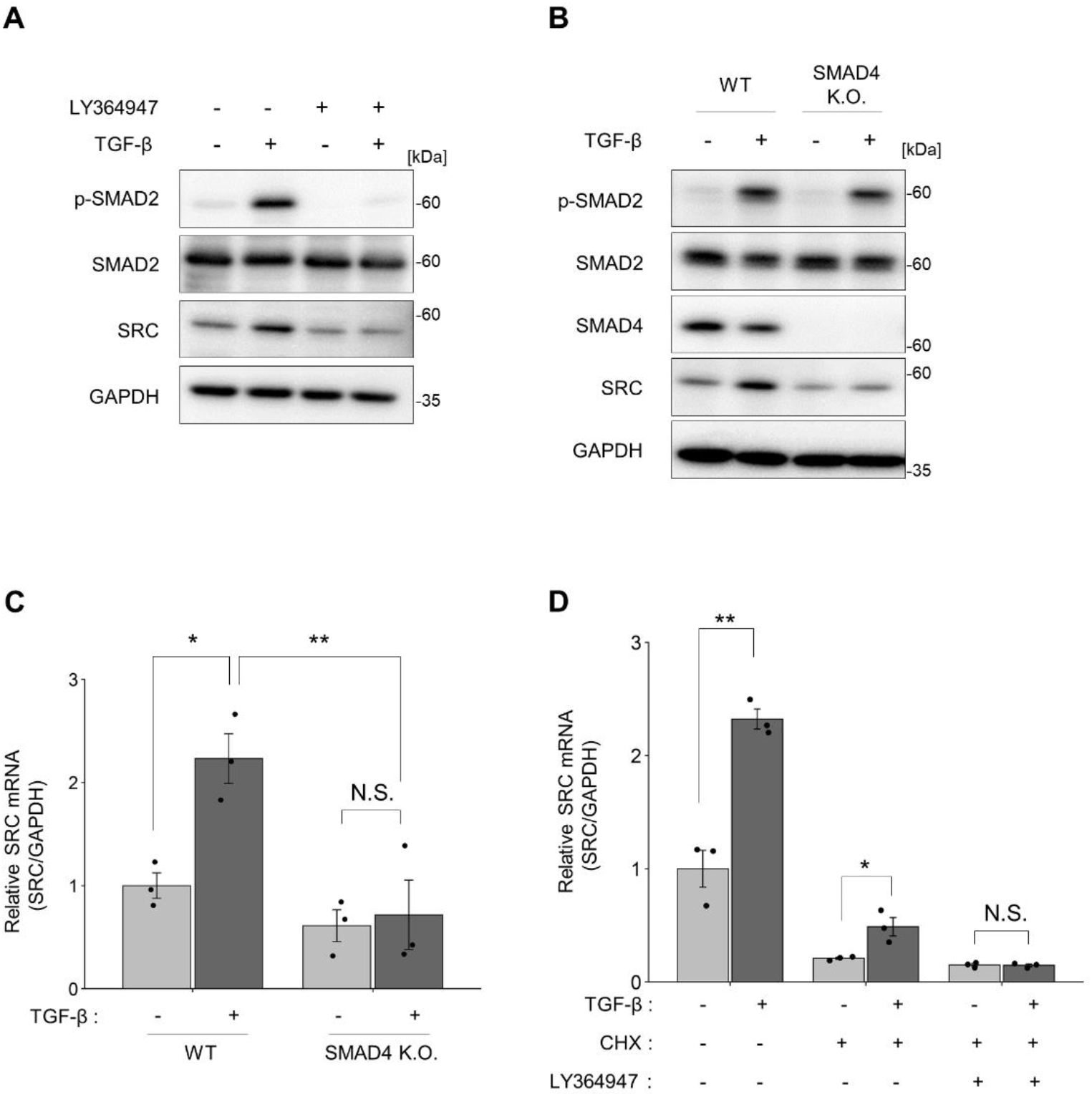
SRC expression is directly regulated by TGF-β/SMAD pathway. (A) MCF10A cells were treated with or without TGF-β1 (10 ng/ml) and the TGFβRI inhibitor LY364947 (2 μg/ml) for 24h. Cell lysates were subjected to immunoblotting using the indicated antibodies. (B, C) Wildtype and SMAD4-knockout clones were treated with TGF-β1 (10 ng/ml) for 24h. (B) Cell lysates were subjected to immunoblotting using the indicated antibodies. (C) Total RNA was isolated and subjected to qPCR. (D) MCF10A cells were pretreated with cycloheximide (10 μg/ml) for 2h, and then treated with or without TGF-β1(10 ng/ml) and TGFβRI inhibitor LY364947 (2 μg/ml) for 8h. Total RNA was isolated and subjected to quantitative real-time polymerase chain reaction PCR. (C and D) The mean ratios ± SEs were obtained from three independent experiments. *, *p* < 0.05; **, *p* < 0.01; N.S., not significantly different; One-way ANOVA with Tukey’s post hoc test in (C); Unpaired two-tailed *t*-test in (D).

### *SRC* intragenic enhancer upregulates *SRC* promoter activity by TGF-β stimulation

We then attempted to identify the genomic region that contributes to TGF-β-induced SRC transcription by chromatin immunoprecipitation (ChIP) with H3K4me3, H3K27Ac, SMAD2, or SMAD3 antibodies. ChIP-Seq results showed that recruitment of SMAD2 and SMAD3 to three regions ~15 kb upstream and downstream of the *SRC* promoter (Fig. 3A and Fig. S2A). Moreover, these regions are marked by H3K27Ac, an epigenetic modification frequently observed in active enhancer sites (Gaarenstroom and Hill, 2014). We refer to these possible regulatory regions as enhancers A, B, and C, respectively, from the upstream *SRC* loci. To assess the contribution of these regions to TGF-β-induced *SRC* promoter activation, we measured the activity of these enhancers using a luciferase reporter assay. The results showed that enhancer B had the ability to upregulate *SRC* promoter activity following TGF-β stimulation (Fig. 3B). Next, we conducted motif analysis using JASPAR database(Castro-Mondragon et al., 2022) and found that there are two putative Smad binding sites (SBS#1 and SBS#2) in enhancer B. To confirm the TGF-β-responsive ability of enhancer B, we introduced mutations into these SBSs by 1 bp substitution or truncation so that they are not predicted as the Smad binging site (Fig. 3C). These mutations reduced TGF-β-induced *SRC* promoter activity, even when another SBS was intact (Fig. 3D and Fig. S2B). These data suggest that the two SBSs in enhancer B are required for upregulation of *SRC* promoter activity. Hereafter, we refer to enhancer B as <u>T</u>GF-β-responsive <u>S</u>RC <u>e</u>nhancer (TSE).

**Figure 3.**
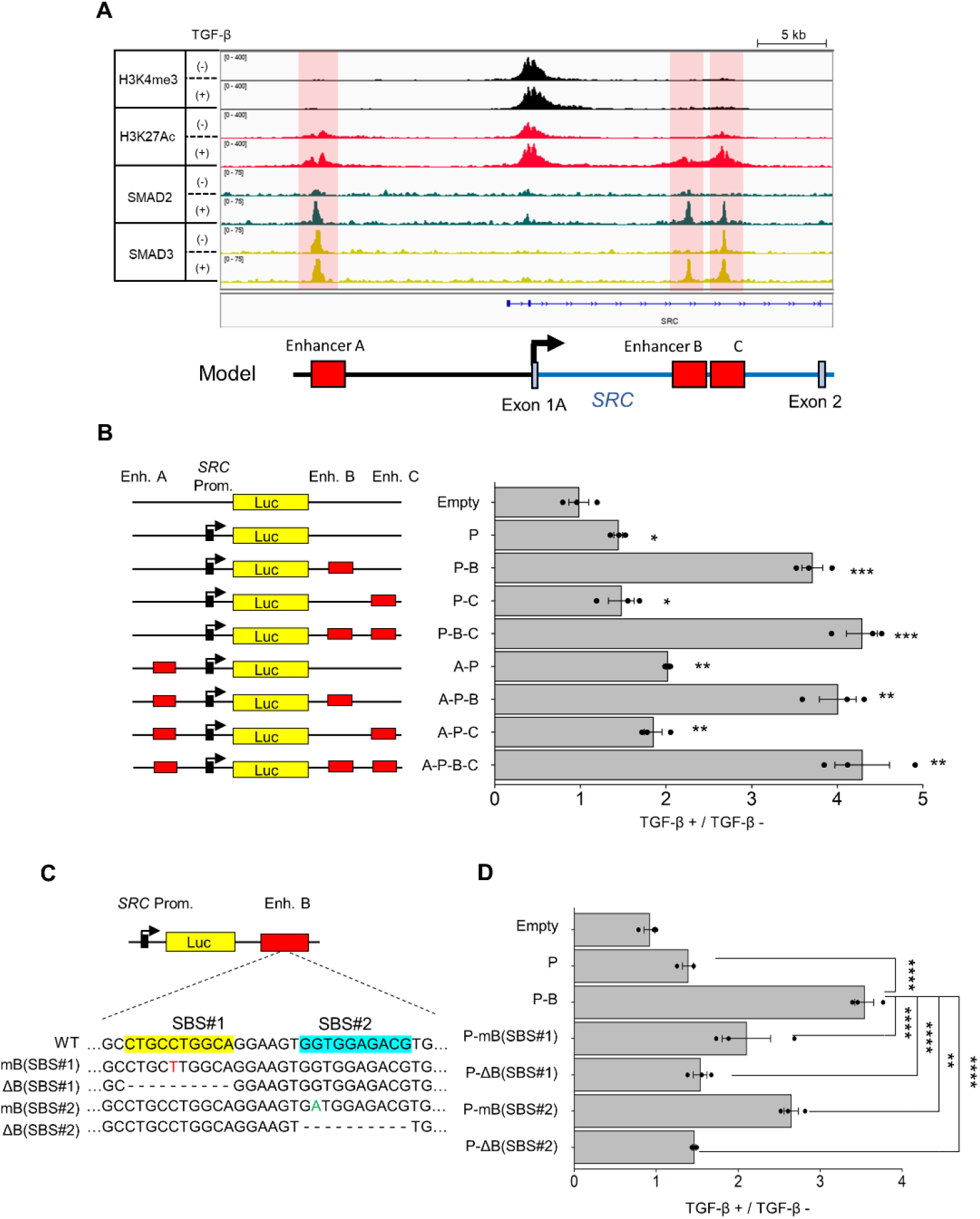
TGF-β-responsive SRC intragenic enhancer increases *SRC* promoter activity. (A) Genomic loci of *the SRC* promoter region. The IP targets are H3K4me3 (black), H3K27Ac (red), SMAD2 (green), and SMAD3(yellow). The ChIP-Seq results were visualized using IGV. (B) Schematic diagram of the F-luciferase reporter vectors is shown on the left, and the result of luciferase reporter assay is shown on the right. The reporter vectors and internal control vector were co-transfected in MCF10A cells overnight, and then transfected cells were treated with TGF-β1(10 ng/ml) for 24h. Cell lysates were subjected to a luciferase reporter assay. Each data point was normalized to the luminescence of the untreated samples. (C) Schematic diagram of SBS-mutant vector constructs. (D) Luciferase reporter assay was conducted using mutant vectors in MCF10A cells. Each data point was normalized to the luminescence of the untreated samples. (B and D) The mean ratios ± SEs were obtained from three independent experiments. *, *p* < 0.05; **, *p* < 0.01; ***, *p* < 0.001; ****, *p* < 0.0001; Unpaired two-tailed *t*-test in (B); One-way ANOVA with Tukey’s post hoc test in (D).

### *SRC* promoter activity is also regulated by the TGF-β/JNK/JUN axis

In addition to SMAD, various transcription factors are required for TGF-β-induced gene expression (Massagué, 2012). Previous studies have shown that the cooperation of the AP-1 transcription factor, which consists of JUN and FOS family proteins, with SMAD plays a pivotal role in breast cancer invasion (Sundqvist et al., 2018; Sundqvist et al., 2020). TGF-β stimulation activates JUN by phosphorylating serine residues via the TGFβ/JNK axis. Moreover, the JUN family proteins directly interact with SMAD proteins to regulate downstream gene expression (Liberati et al., 1999). Consistent with these studies, motif enrichment analysis using ChIP-Seq results for SMAD2 in the presence of TGF-β suggested that AP-1-related motifs were included in the SMAD complex binding site (Fig. S3A). Furthermore, overlap analysis revealed that over 70% of SMAD complex binding regions overlapped with JUN binding regions (Fig. S3B). ChIP-Seq analysis also revealed the co-localization of SMAD and JUN in the TSE (Fig. 4A). Furthermore, the luciferase reporter assay showed that TGF-β-induced SRC promoter activity was decreased by mutation of the AP-1 binding site in TSE (Fig. 4B).

**Figure 4.**
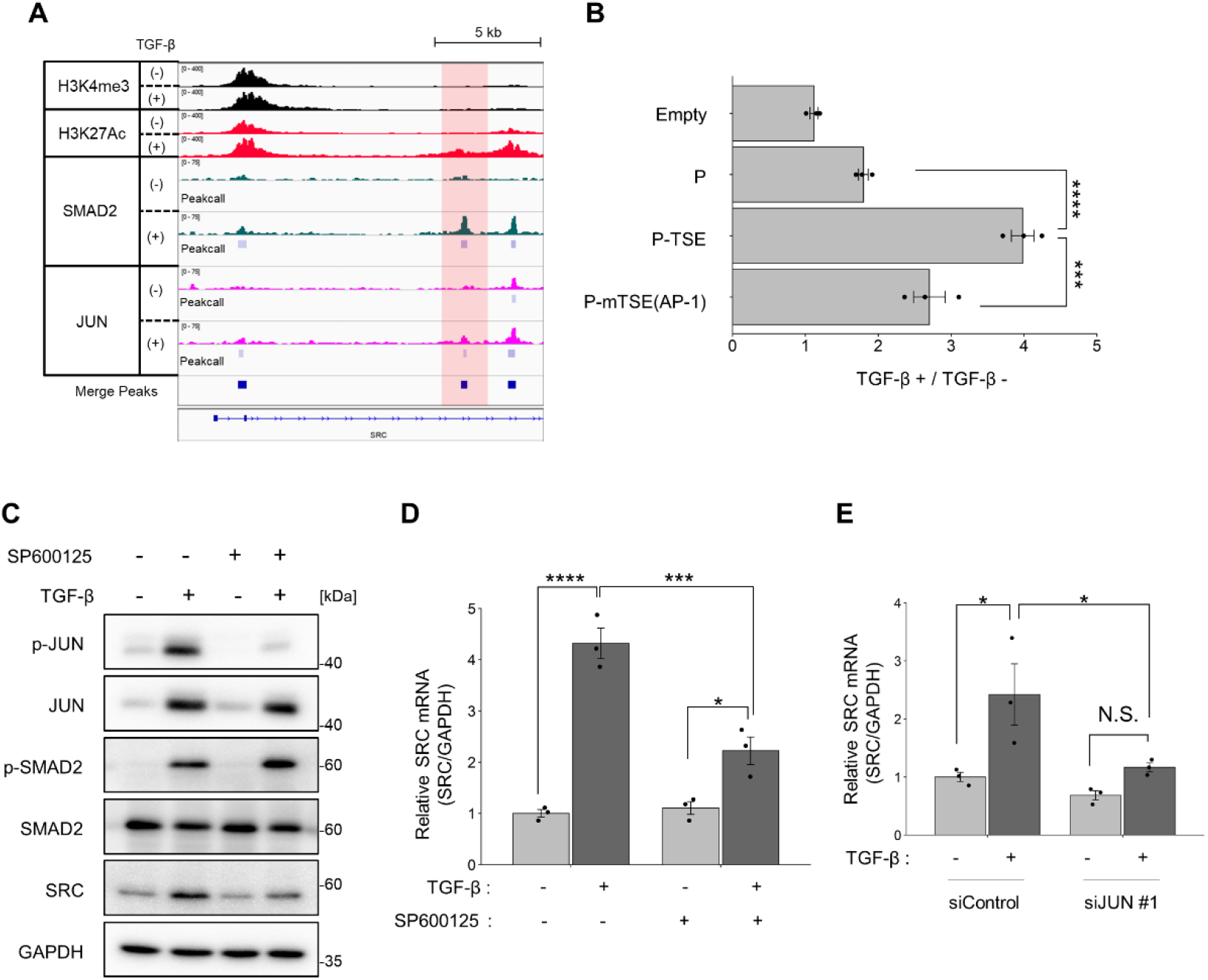
JUN is required for TSE-mediated SRC promoter activation. (A) Genomic loci of *the SRC* promoter region. The IP targets are H3K4me3 (black), H3K27Ac (red), SMAD2 (green), and JUN (magenta). The ChIP-Seq results were visualized using IGV. Peak call analysis was conducted using the findPeaks program in HOMER. (B) The result of luciferase reporter assay. Reporter vectors and internal control vector co-transfected in MCF10A overnight, and transfected cells were treated with TGF-β1(10 ng/ml) for 24h. Cell lysates were subjected to a luciferase reporter assay. Each data point was normalized to the luminescence of the untreated samples. (C, D) MCF10A cells were treated with or without TGF-β1 (10 ng/ml) and SP600125 (20 μM) for 24h. (C) Cell lysates were subjected to immunoblotting using the indicated antibodies. (D) Total RNA was isolated and subjected to qPCR analysis. (E) MCF10A cells were treated with the indicated siRNAs overnight and then with or without TGF-β1(10 ng/ml) for 24h. Total RNA was isolated and subjected to quantitative real-time polymerase chain reaction PCR. (B, D, and E) The mean ratios ± SEs were obtained from three independent experiments. *, *p* < 0.05; ***, *p* < 0.001; ****, *p* < 0.0001; One-way ANOVA with Tukey’s post hoc test.

To further verify the involvement of the JNK/JUN signaling pathway in the regulation of SRC transcription, we examined the effects of JNK inhibitor treatment and JUN protein knockdown. TGF-*β* stimulation induced the expression and the phosphorylation of JUN protein, and the JNK inhibitor SP600125 inhibited the JNK-mediated phosphorylation of JUN and reduced TGF-β-induced SRC expression (Fig. 4C and D). Moreover, JUN knockdown reduced TGF-β-induced SRC expression (Fig. 4E, Fig. S3C and D). These results demonstrated that *SRC* promoter activity is synergistically regulated by the TGF-β/JNK/JUN axis.

### TSE is essential for TGF-β-induced SRC activation; however, EMT-associated morphological changes occur even in the absence of TSE

To analyze the effects of TGF-β-induced SRC expression on cellular functions, we established TSE-mutant MCF10A cell lines using the CRISPR/Cas9 system (Zientek-Targosz et al., 2008) (Fig. S4). In these cell lines, TGF-β-induced SRC expression was completely suppressed at both mRNA and protein levels (Fig. 5A-C). These results verified that TSE-mediated SRC upregulation occurs via endogenous SRC transcriptional machineries. Notably, although the total amount of active SRC (pY419) was increased by TGF-β stimulation in wild-type cells (Fig. 5B), the ratio of active SRC to total SRC did not change in either wild-type or TSE-mutant cells (Fig. 5D). These results suggested that TGF-β-induced SRC activation is mainly attributable to the upregulation of TSE-mediated SRC expression.

**Figure 5.**
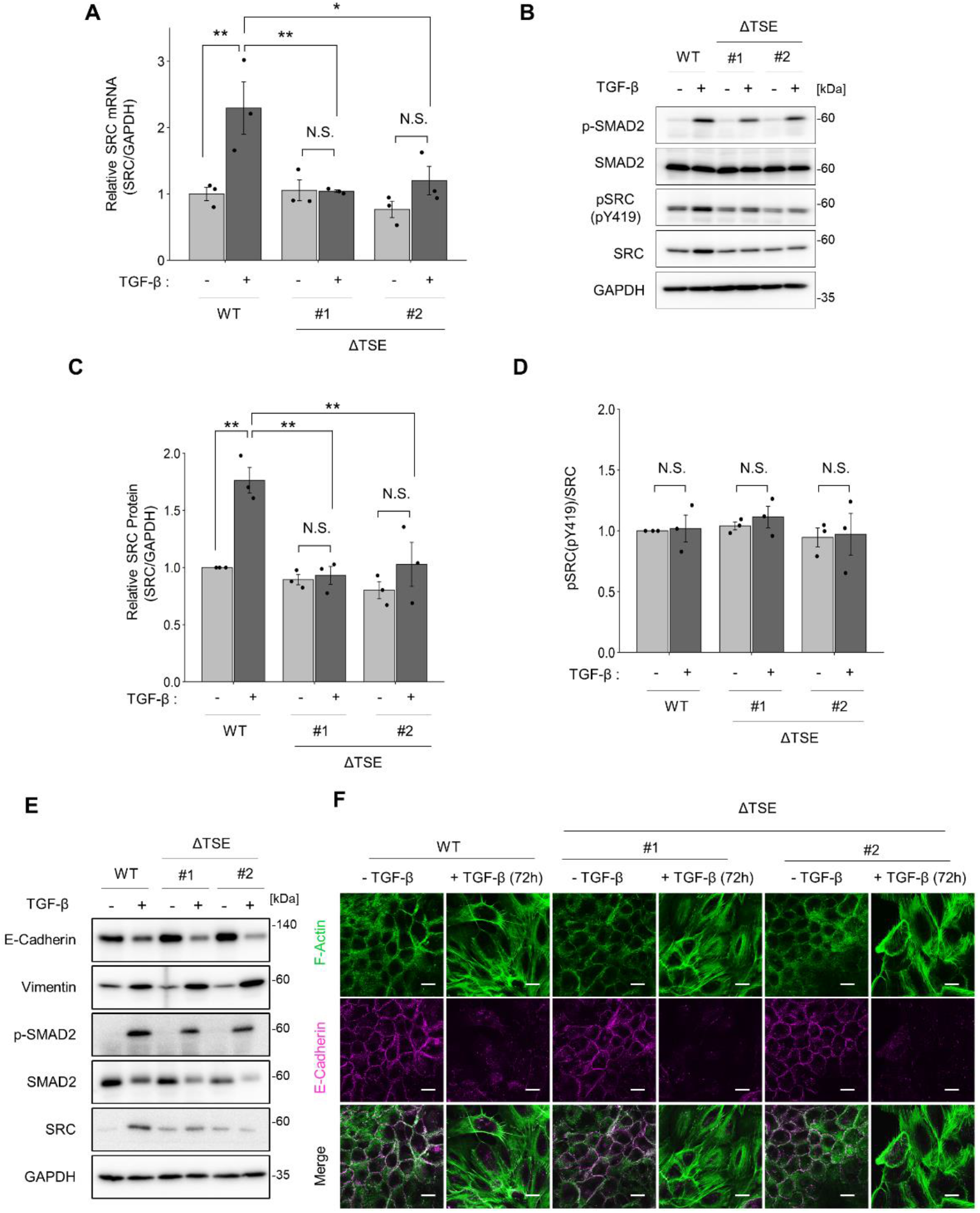
TGF-β stimulation induces EMT-associated morphological changes in TSE-deficient cell lines. (A, B) wild-type MCF10A cells and ΔTSE clones were treated with TGF-β1(10 ng/ml) for 24h. (A) Total RNA was isolated and subjected to qPCR. (B) Cell lysates were subjected to immunoblotting using the indicated antibodies. (C, D) Quantification of (C) SRC protein levels and (D) SRC-pY419 in the immunoblot analysis shown in (B). (E, F) wild-type MCF10A cells and ΔTSE clones were treated with TGF-β1(10 ng/ml) for 72h. (E) Cell lysates were subjected to immunoblotting using the indicated antibodies. (F) Cells were subjected to immunofluorescence staining for F-actin and E-cadherin expression. The scale bar represents 10 μm. (A, C, and D) The mean ratios ± SEs were obtained from three independent experiments. *, *p* < 0.05; **, *p* < 0.01; N.S., not significantly different; One-way ANOVA with Tukey’s post hoc test.

Previous studies have shown that constitutively active SRC (e.g., v-Src, SrcY527F mutant) promotes the disorganization of cell-cell contacts by directly phosphorylating E-cadherin and activating the SRC/FAK/ERK/MLCK/myosin pathway, resulting in the induction of EMT (Avizienyte and Frame, 2005; Avizienyte et al., 2004; Fujita et al., 2002; Webb et al., 2004). However, another study revealed that endogenous SRC activation was not necessary for TGF-β-induced EMT (Maeda et al., 2006). To elucidate the role of SRC in EMT, we evaluated the effects of TGF-β-induced SRC activation on the EMT process using the TSE mutant MCF10A cell line. Immunoblot analysis showed that the TSE mutation did not significantly affect the repression of E-cadherin and induction of vimentin expression (Fig. 5E). In addition, EMT-associated cellular traits, including the reduction in E-cadherin-mediated cell-cell adhesion and the formation of stress fibers, were observed in both wild-type and TSE mutant cell lines (Fig. 5F). Taken together, it is likely that TGF-β-induced SRC activation at the endogenous level and EMT-associated morphological changes are independent events, at least in MCF10A cells.

### TSE regulates EMT-associated cell motility by upregulating SRC-mediated FAK phosphorylation

Finally, we investigated the effects of TSE-mediated SRC upregulation on EMT. The cell proliferation assay showed that the growth arrest effect was observed at the same ratio in both wild-type and TSE-mutant cell lines, implying that TSE-mediated SRC expression does not significantly affect cell proliferation (Fig. S5). Since the total amount of active SRC increased (Fig. 5B), we tried to identify the proteins phosphorylated by SRC. Immunoblot analysis using an anti-phospho-tyrosine antibody (pY1000) showed that phosphorylation of ~130 kDa proteins was increased by TGF-β stimulation in the wild-type cell line, but not in TSE mutant cell lines (Fig. 6A). This result suggests that TSE-mediated SRC upregulation results in phosphorylation of these proteins. We examined several tyrosine-phosphorylated proteins previously reported as SRC substrates (Guarino, 2010) and found that SRC-mediated phosphorylation of FAK was significantly decreased in TSE mutant cell lines (Fig. 6B-D). Importantly, the FAK inhibitor Y15, which selectively interacts with Y397 of FAK and inhibits autophosphorylation of the residue (Golubovskaya et al., 2008), suppressed SRC-Y419 phosphorylation and subsequent FAK-Y576/577 phosphorylation (Westhoff et al., 2004) (Fig. 6E). This result suggests that TGF-β-induced SRC expression promotes formation and activation of the SRC/FAK circuit. Because SRC/FAK signaling plays a central role in cell migration (Avizienyte and Frame, 2005; Guarino, 2010; Le Coq et al., 2022; Westhoff et al., 2004), we examined the effects of TSE-mediated SRC upregulation on cell motility using a wound healing assay. The results showed that TGF-β-induced cell migration was decreased in the TSE mutant cell lines (Fig. 6F and G). These results indicated that TSE-mediated SRC upregulation is crucial for FAK activation to maximize TGF-β-induced cell migration.

**Figure 6.**
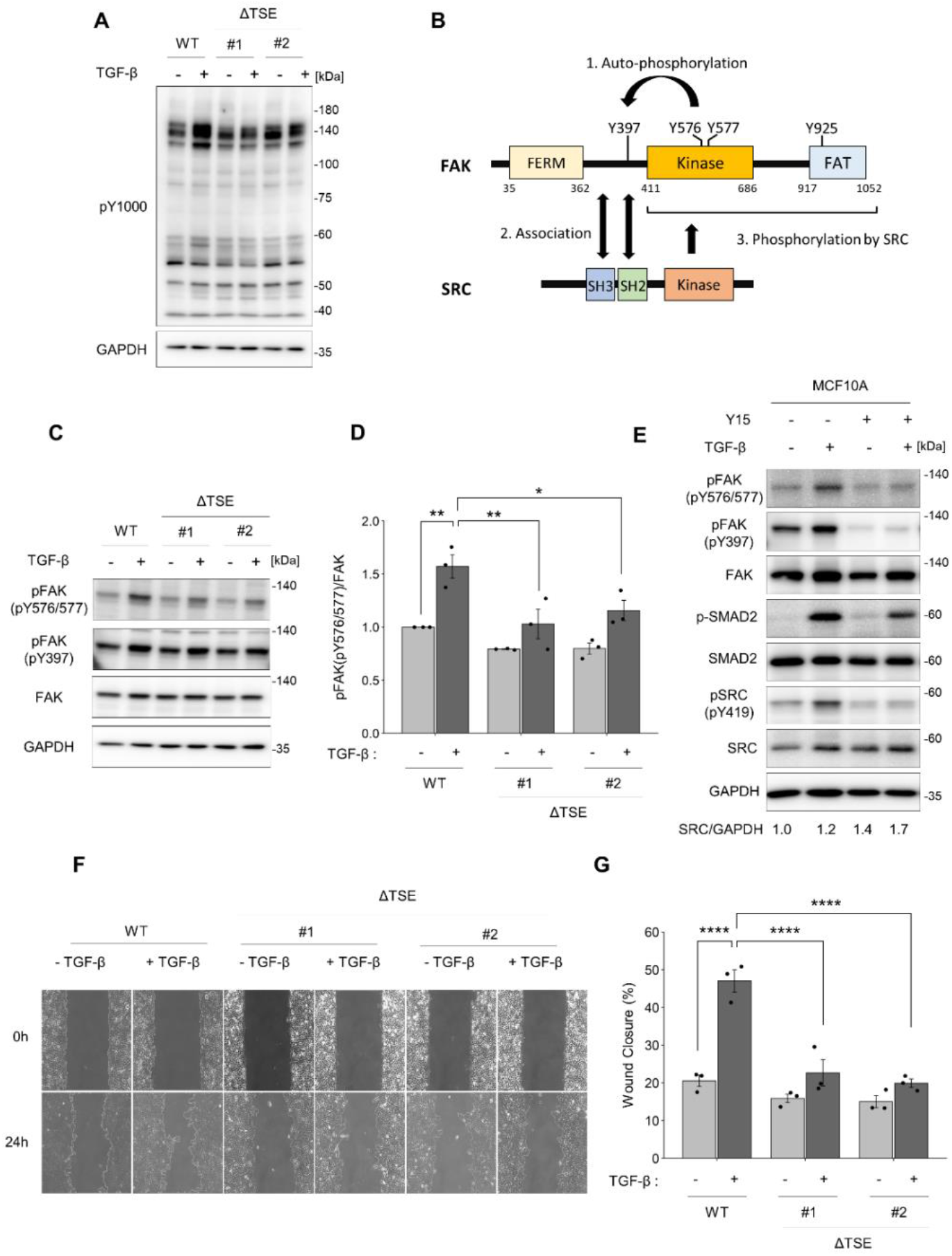
Deletion of TSE attenuates the SRC-mediated FAK phosphorylation and EMT-associated cell motility. (A) wild-type MCF10A cells and ΔTSE clones were treated with TGF-β1(10 ng/ml) for 24h. Cell lysates were subjected to immunoblotting using the indicated antibodies. (B) Schematic diagram of the interactions between FAK and SRC. (C) wild-type MCF10A cells and ΔTSE clones were treated with TGF-β1(10 ng/ml) for 24h. Cell lysates were subjected to immunoblotting using the indicated antibodies. (D) Quantification of FAK-pY576/577 level in immunoblot analysis shown in (C). (E) MCF10A cells were treated with or without TGF-β1(10 ng/ml) and the FAK inhibitor Y15 (5 μM) for 24h. Cell lysates were subjected to immunoblotting using the indicated antibodies. (F) Wound healing assay of wild-type MCF10A cells and ΔTSE clones treated with or without TGF-β1 (10 ng/ml) for 24h. (G) Quantification of the wound closure rate using the images shown in (F). (D and G) The mean ratios ± SEs were obtained from three independent experiments. *, *p* < 0.05; **, *p* < 0.01; ****, *p* < 0.0001; One-way ANOVA with Tukey’s post hoc test.

## Discussion

We investigated the mechanisms underlying TGF-β-induced upregulation of SRC in MCF10A cells and identified a TGF-β-responsive SRC enhancer (TSE) that induces the transcriptional activation of the *SRC* promoter via the TGF-β/SMAD and TGF-β/JNK/JUN signaling pathways. Because SRC has oncogenic potential, its expression and activity are strictly regulated by post-transcriptional and post-translational pathways. In post-transcriptional regulation, miRNAs directly bind to SRC mRNA, resulting in the degradation and inhibition of SRC mRNA translation (Majid et al., 2011; Okuzaki et al., 2020; Oneyama and Okada, 2015; Wang et al., 2014). In post-transcriptional regulation, the specific activity of SRC is regulated by C-terminal regulatory phosphorylation (Nada et al., 1991; Okada, 2012), and its protein level is regulated by degradation via the lysosome (Tsai et al., 2019) and proteasome (Kim et al., 2004; Mukhopadhyay et al., 2016) systems or by excretion via small extracellular vesicles (Tanaka et al., 2020). In this study, we focused on the transcriptional regulatory systems of *SRC*. This is the first report to demonstrate the transcriptional regulation of *SRC* induced by extracellular stimulation.

According to the COSMIC database, SRC is overexpressed in various human cancers; however, mutations in *SRC* are rare (Fig. S6A). Additionally, The Cancer Genome Atlas (TCGA) database analysis revealed that SRC is significantly overexpressed in human breast cancer (Fig. S6B). In colorectal and pancreatic cancers, the TGF-β signaling pathway is frequently abolished by mutations in *SMAD4* or *TGFBR2* (Joan Massagué, 2008; Levy and Hill, 2006). In contrast, TGF-β/SMAD signaling machinery is preserved in breast cancer (Joan Massagué, 2008). Furthermore, TGF-β signaling plays a pivotal role in promoting tumor cell invasion and bone metastasis in breast cancer (Deckers et al., 2006; Sundqvist et al., 2018; Sundqvist et al., 2020). These studies, together with our findings, provide a possible link between the upregulation of TGF-β signaling and SRC, particularly in breast cancer.

We previously reported that TGF-β2 stimulation induces SRC activation by upregulating the SRC scaffolding protein CASL, without upregulating SRC expression in primary human trabecular meshwork cells (Tsukamoto et al., 2019). Therefore, the cell-type specificity that determines whether TGF-β can induce SRC expression needs to be further elucidated. In this study, we showed that the AP-1 family protein JUN, in addition to SMAD, binds to the TSE, resulting in *SR*C promoter activation. AP-1, in combination with EGF signaling, plays an essential role in regulating the transcriptional program of SMAD and promotes TGF-β-induced invasiveness by cooperating with SMAD in ΔNp63-expressing breast cancer cell lines (Sundqvist et al., 2020). Consistent with this study, we found that co-stimulation with TGF-β and EGF induced SRC expression in normal human epidermal keratinocytes (NHEK) expressing ΔNp63 (Fig. S7A-C). These results imply that the cooperative mechanisms of SMAD and AP-1 may determine the cell-type specificity of TGF-β-induced SRC expression. In addition, TSE contains epigenetic modifications of H3K4me1 without H3K27Ac, which represents the poised enhancer, in the basal state of MCF10A and NHEK (Fig. S7D). This result suggests that cell-type-specific epigenetic states may also reveal cell-type specificity.

We further found that TSE-mediated SRC expression was crucial for increased FAK phosphorylation. SRC/FAK signaling accelerates tumor invasiveness by regulating integrin-mediated cell adhesion, cell motility, and extracellular matrix degradation. Therefore, TSE-mediated activation of the SRC/FAK circuit may promote EMT-associated cancer cell migration and invasion. As SRC is a key protein in tumor progression, various small-molecule inhibitors of SRC have been developed for anti-cancer therapy (Mayer and Krop, 2010). However, ATP-competitive SRC inhibitors induce a conformational change in SRC, leading to stable interaction with FAK at focal adhesions. Subsequent reduction in inhibitor concentration enables SRC to readily phosphorylate FAK and paradoxically activate the FAK-Grb2-Erk signaling pathway (Higuchi et al., 2021). In this study, we demonstrated that the inhibition of TGF-β-induced SRC expression results in the suppression of SRC activity and EMT-associated cell migration. This result suggests that the machinery of TSE-mediated SRC transcription can be a novel inhibitory target of SRC without causing unexpected effects such as conformational changes in SRC.

In summary, we demonstrated that a TGF-β-responsive enhancer upregulates SRC expression and that TGF-β-induced SRC expression is essential for EMT-associated cell migration. These findings raise the possibility that the overexpression of SRC observed in a subset of human cancers is induced by TGF-β secreted in the tumor microenvironment. Further investigation of the transcriptional mechanisms of *SRC* may provide promising targets for anticancer drugs that prevent tumor invasion and metastasis.

## Experimental procedures

### Cell culture

MCF10A cells were cultured in DMEM/F12 (Nacalai Tesque) supplemented with 5% horse serum (Gibco), 20 ng/ml EGF (PeproTech), 0.5 μg/ml hydrocortisone (Sigma), 10 μg/ml insulin (Wako), 100 ng/ml Cholera toxin (Bio Academia), and a penicillin–streptomycin solution (Nacalai Tesque). BT-549 cells were cultured in RPMI1640 (Nacalai Tesque) containing 10% FBS, 0.023 U/ml insulin (Wako), and penicillin–streptomycin solution (Nacalai Tesque). Normal Human Epidermal Keratinocyte (NHEK) cells were purchased from PromoCell and cultured in keratinocyte growth medium 2(PromoCell). The cells were maintained in a CO_2_ incubator at 37 °C and 5% CO_2_. Recombinant Human TGF-β1 (CHO derived) was purchased from PeproTech.

### Antibodies and inhibitors

The following primary antibodies were used in this study: anti-v-Src (OP07) antibody was purchased from Sigma-Aldrich. Anti-phospho-Src family (pY416) (2101), anti-phospho-tyrosine (8954), anti-Smad2 (5339), anti-phospho-Smad2 (3108), anti-Smad3 (9523), anti-c-Jun (9165), anti-p-c-Jun (3270), anti-vimentin (5741), anti-p63-α (13109), anti-H3K4me3 (9751), and anti-H3K27c (8173) were purchased from Cell Signaling Technology. Anti-E-cadherin (610181), anti-N-cadherin (610920), and anti-FAK (610087) antibodies were purchased from BD Biosciences. Anti-GAPDH (32233), anti-SP1 (59), anti-Smad4 (7966), and anti-phospho-FAK (pY576/577) (21831) antibodies were purchased from Santa Cruz Biotechnology. Anti-Claudin-1 (717800) and anti-phospho-FAK (pY397) (700255) antibodies were purchased from Thermo Fisher Scientific.

The following inhibitors were used in this study: TGFβR-I inhibitor LY364947 (S2805), JNK inhibitor SP600125 (S1460) and FAK inhibitor Y15 (S5321) were purchased from Selleck. Mitomycin C Solution (20898-21) was purchased from Nacalai Tesque. Actinomycin D (018-21264) was purchased from Wako.

### Plasmid construction and gene transfer

DNA sequences of Enhancer A, B, and C were cloned by PCR using human genomic DNA as the template and subcloned into the pGV-B2 plasmid (Toyo Bnet). TSE mutants were generated by mutagenesis PCR. Plasmids were transfected with PEI MAX (Polysciences Inc). Primers used for this experiment are listed in Table S1. siRNAs were purchased from Sigma-Aldrich and transfected with Lipofectamine RNAiMAX (Thermo Fisher Scientific). siRNAs used are listed in Table S2.

### Immunoblotting

Cells were lysed with RIPA buffer (50 mM Tris-HCl [pH 7.4], 150 mM NaCl, 1% NP-40, 0.1% SDS, 1 mM EDTA, 0.5% sodium deoxycholate, 1 mM Na3VO4, 20 mM NaF, 1 mM PMSF, and protease inhibitor cocktail (Nacalai Tesque)) and debris was removed by centrifugation at 15,000×g for 10 min. HRP-conjugated anti-mouse or anti-rabbit IgG (Zymed Laboratories Inc.) was used as secondary antibody. All immunoblots were visualized using a Luminograph II System (Atto) and quantified using the ImageJ software.

### Immunofluorescence microscopy

Cells were grown on type I collagen-coated coverslips, fixed with 4% paraformaldehyde, and permeabilized with PBS containing 0.03% Triton X-100. The permeabilized cells were blocked with Blocking One (Nacalai Tesque) and incubated with primary antibodies overnight, followed by incubation with Alexa Fluor 488-conjugated phalloidin and Alexa Fluor 594-conjugated secondary antibodies. Immunostained samples were observed under an FV1000 confocal microscope (Olympus).

### RNA isolation and qRT-PCR

Total RNA was isolated from the cells using the NucleoSpin RNA kit (Macherey-Nagel), and cDNA was prepared using ReverTra Ace RT Master Mix (TOYOBO) according to the manufacturer’s instructions. Real-time PCR was performed using the QuantStudio 5 Real-Time PCR System (Applied Biosystems) and the THUNDERBIRD NEXT SYBR qPCR Mix (TOYOBO). Gene expression levels were normalized to GAPDH expression. Primers used for this analysis are listed in Table S3.

### CRISPR/Cas9-based gene knockout and mutagenesis

Sequences of gRNA were designed using CRISPRdirect (DBCLS) and inserted into pX458. Target-gRNA contained pX458 were transfected into MCF10A cells with PEI MAX (Polysciences Inc). The next day, EGFP-positive cells were isolated using a FACSAria III sorter (BD Biosciences). Genomic mutations were confirmed by immunoblotting and sequencing. The sequences of the gRNA and primers used for genotyping are listed in Table S4.

### ChIP-Sequencing and data processing

Cells were cultured in a 15 cm dish to 80–90% confluence (approximately 2×10^7^ cells), and one dish was used for immunoprecipitation. ChIP experiments were performed using the SimpleChIP Enzymatic Chromatin IP kit (9003, Cell Signaling Technologies), according to the manufacturer’s instructions. The antibodies used for ChIP were anti-H3K4me3 (9751), anti-H3K27c (8173), anti-Smad2 (5339), anti-Smad3 (9523), anti-c-Jun (9165), and anti-JunB (3753) which were purchased from Cell Signaling Technologies.

ChIP-seq libraries were prepared using the KAPA Hyper Preparation Kit. Sequencing was performed on NovaSeq 6000 platform in a 101+101 base paired-end mode. Illumina-TruSeq Adapter Trimming was performed on ChIP-seq reads, using cutadapt v2.7, discarding reads left with < 30 bp. ChIP-seq reads were aligned to the UCSC hg19 genome using Bowtie2 version 2.3.5.1. bigWig files were generated using the bamCoverage software of deepTools package.

Peak call analysis, merge peak analysis and find motif analysis were performed by “findPeaks” program, “mergePeaks” program and “findMotifsgenome.pl” program, respectively in the HOMER version 4.11 package. ChIP-Seq data for H3K4me1 in MCF10A cells were obtained from GSE85158. ChIP-Seq data for SMAD2/3 in BT-549 cells were obtained from GSE104352. ChIP-Seq data for H3K4me1 and H3K27Ac in NHEK were obtained from GSE29611.

### Luciferase Reporter Assay

For cell preparation, 5×10^4^ cells were seeded in 48-well plates and cultured for 24h. Cells were co-transfected with the pGV-B2 reporter vector and pRL-TK internal control vector for 20h. The following day, cells were stimulated with TGF-β for 24h by refreshing the medium with or without 10 ng/ml TGF-β, and luciferase activity was assayed using a PicaGene Dual Sea Pansy Luminescence Kit (Toyo Bnet) according to the manufacturer’s instructions.

### Wound Healing Assay

For cell preparation, 3×10^4^ cells were seeded into 2-well silicone culture insert (ib80209, ibidi) in a 35 mm dish and cultured overnight. To stop cell proliferation, cells were treated with 10 μg/ml Mitomycin C containing medium for 2h, and then the culture inserts were removed. Cells were washed twice with PBS and cultured in a medium with or without 10 ng/ml TGF-β for 24h. Cell pictures were taken at the timepoint of 0h, 24h.

### Statistics

All statistical analyses were performed using R version 4.0.3 and Microsoft Excel. One-way analysis of variance (ANOVA) with Tukey’s post hoc test was used for multiple group comparisons. Unpaired two-tailed *t*-tests were performed for two groups comparison. A *p*-value less than 0.05 was considered significant. All data were derived from at least three independent experiments.

## Data availability

Access to raw data concerning this study was submitted under Gene Expression Omnibus (GEO) accession number GSE216432.

## Supporting information

This article contains supporting information.

## Acknowledgments

The authors thank Dr. Hiroki Nishida for experimental assistance. We also acknowledge the NGS core facility of the Genome Information Research Center at the Research Institute for Microbial Diseases of Osaka University for their support with DNA sequencing and data analyses. This work was supported by JPSP KAKENHI (Grant numbers 19H03504 and 19H04962 to M.O., and 19K07639 to K.K.) and JST SPRING (Grant Number JPMJSP2138 to S.N.). We would like to thank Editage (www.editage.com) for the English language editing.

## Conflict of interest

The authors declare no conflicts of interest regarding contents of this article.

## Abbreviations

TGF-β: (Transforming growth factor-β)
SFK: (Src Family Kinase)
AP-1: (Activator protein-1)
FAK: (Focal adhesion kinase)
EMT: (Epithelial-mesenchymal transition)
ECM: (Extracellular matrix)
MMP: (Matrix metalloproteinase)
SBS: (SMAD binding site)
TSE: (TGF-β-responsive SRC enhancer)

---

**Figure S1.**
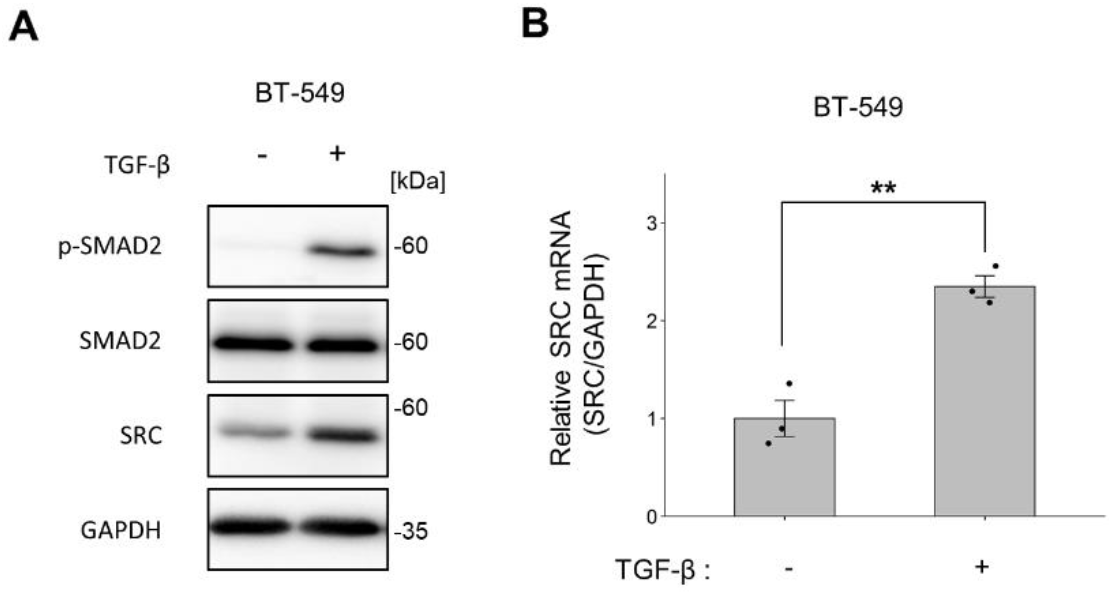
TGF-β stimulation upregulates *SRC* transcription in triple negative breast cancer cell line BT-549. (A, B) triple-negative breast cancer cell line BT-549 was treated with TGF-β1 (10 ng/ml) for 24h. (A) Cell lysates were subjected to immunoblotting using the indicated antibodies. (B) Total RNA was isolated and subjected to qPCR. (B) Mean ratios ± SEs were obtained from three independent experiments. **, *p* < 0.01; Unpaired two-tailed *t*-test.

**Figure S2.**
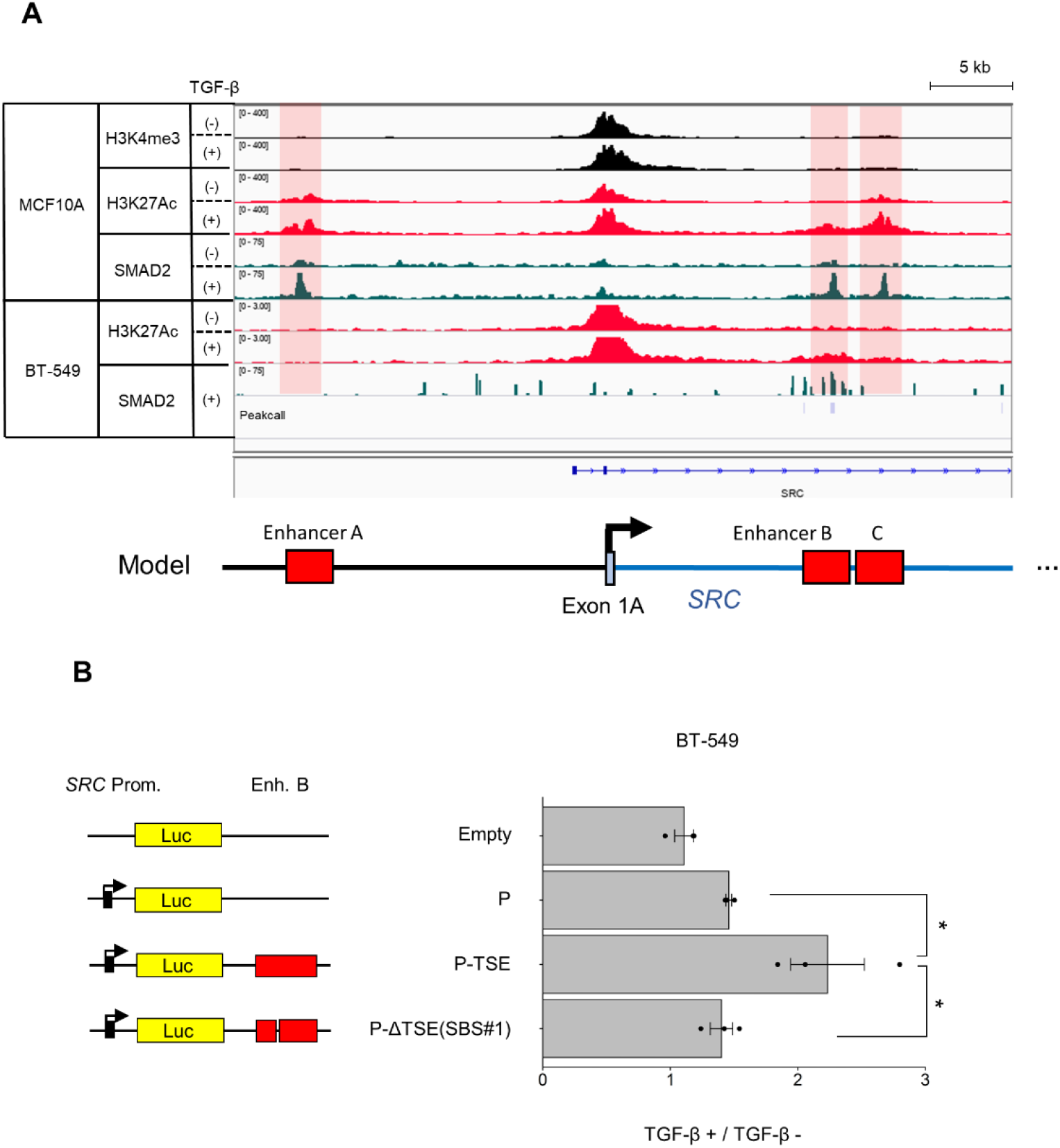
TGF-β-responsive SRC intragenic enhancer increases *SRC* promoter activity in BT-549. (A) Genomic loci of the *SRC* promoter region. The IP targets are H3K4me3 (black), H3K27Ac (red), and SMAD2 (green). The ChIP-Seq results in MCF10A and BT-549 cells were visualized using IGV. Peak call analysis was conducted using the findPeaks program in HOMER. (B) Luciferase reporter assay was conducted using the SBS mutant vector in BT-549 cells. Each data point was normalized to the luminescence of the untreated samples. (B) Mean ratios ± SEs were obtained from three independent experiments. \**p* < 0.05; One-way ANOVA with Tukey’s post hoc test.

**Figure S3.**
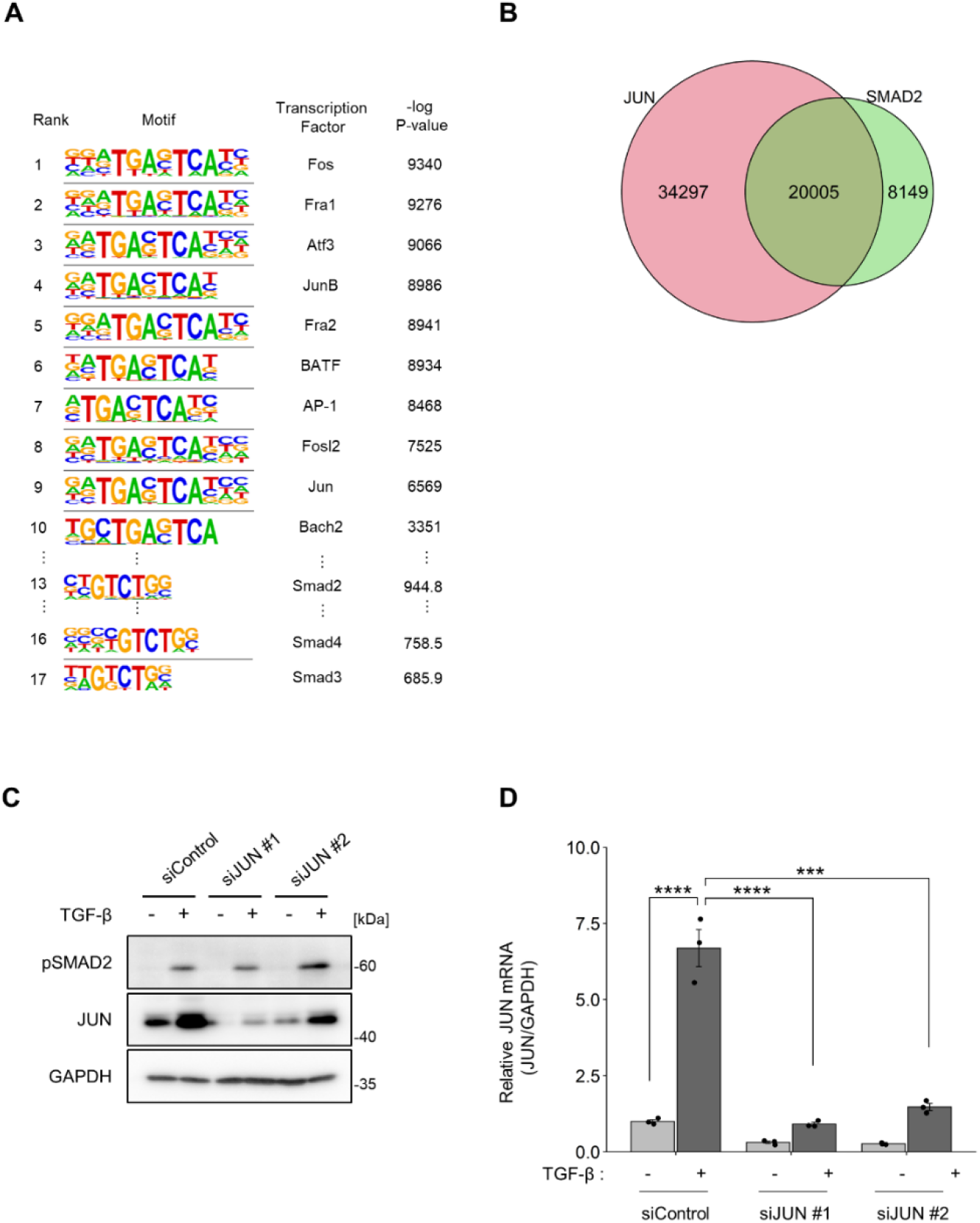
Motif enrichment analysis focusing on JUN, and knock down of JUN by siRNAs in MCF10A. (A) Motifs enriched in the SMAD complex binding site in TGF-β-stimulated MCF10A cells. (B) Comparison of the locus of JUN binding sites and SMAD complex binding sites.(C, D) MCF10A cells were treated with the indicated siRNAs overnight and then with or without TGF-β1(10 ng/ml) for 24h. (C) Cell lysates were subjected to immunoblotting using the indicated antibodies. (D) Total RNA was isolated and subjected to qPCR. (D) Mean ratios ± SEs were obtained from three independent experiments. ***, *p* < 0.001; ****, *p* < 0.0001; One-way ANOVA with Tukey’s post hoc test.

**Figure S4.**
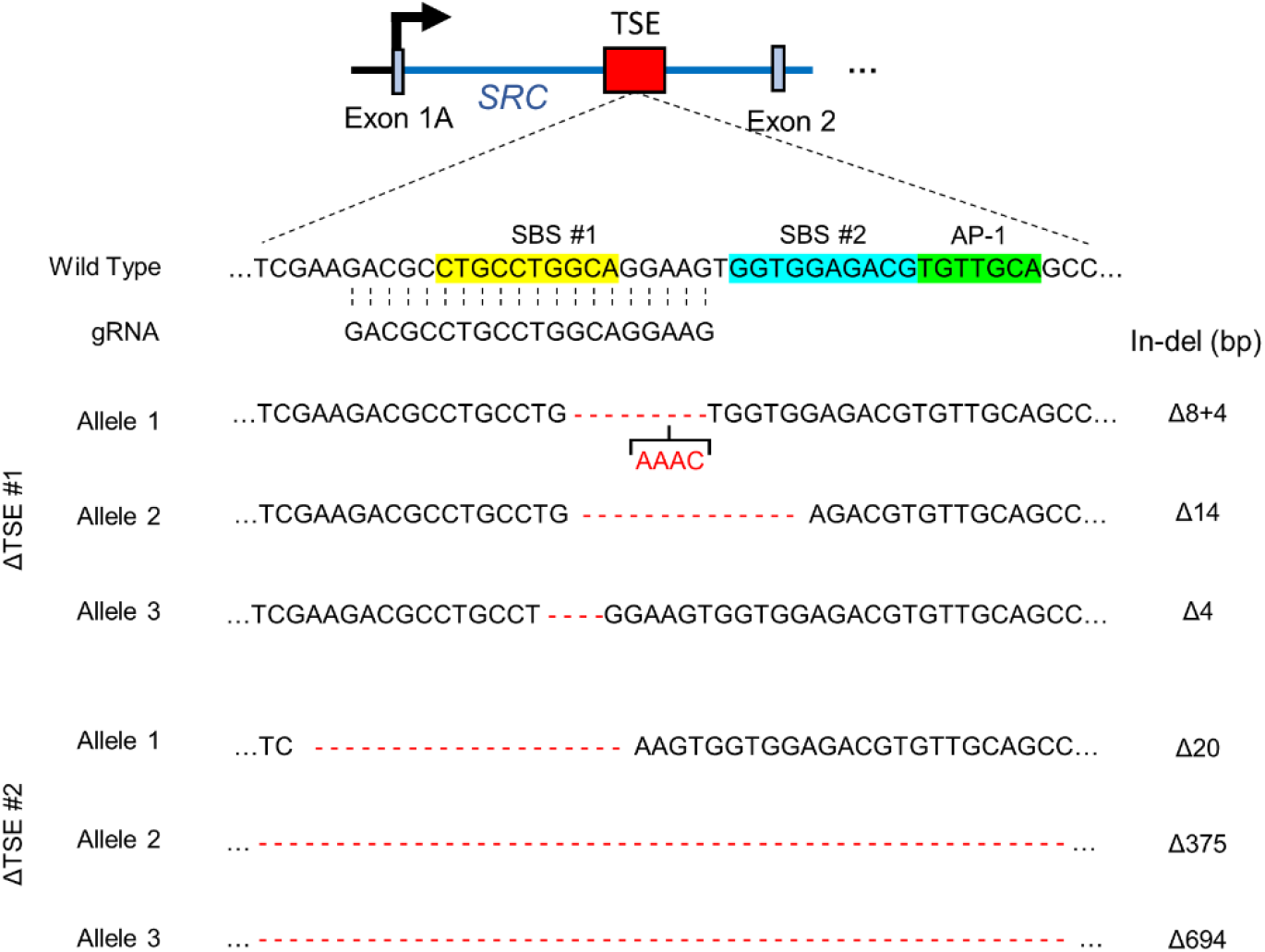
DNA sequences in ΔTSE MCF10A cell lines.

**Figure S5.**
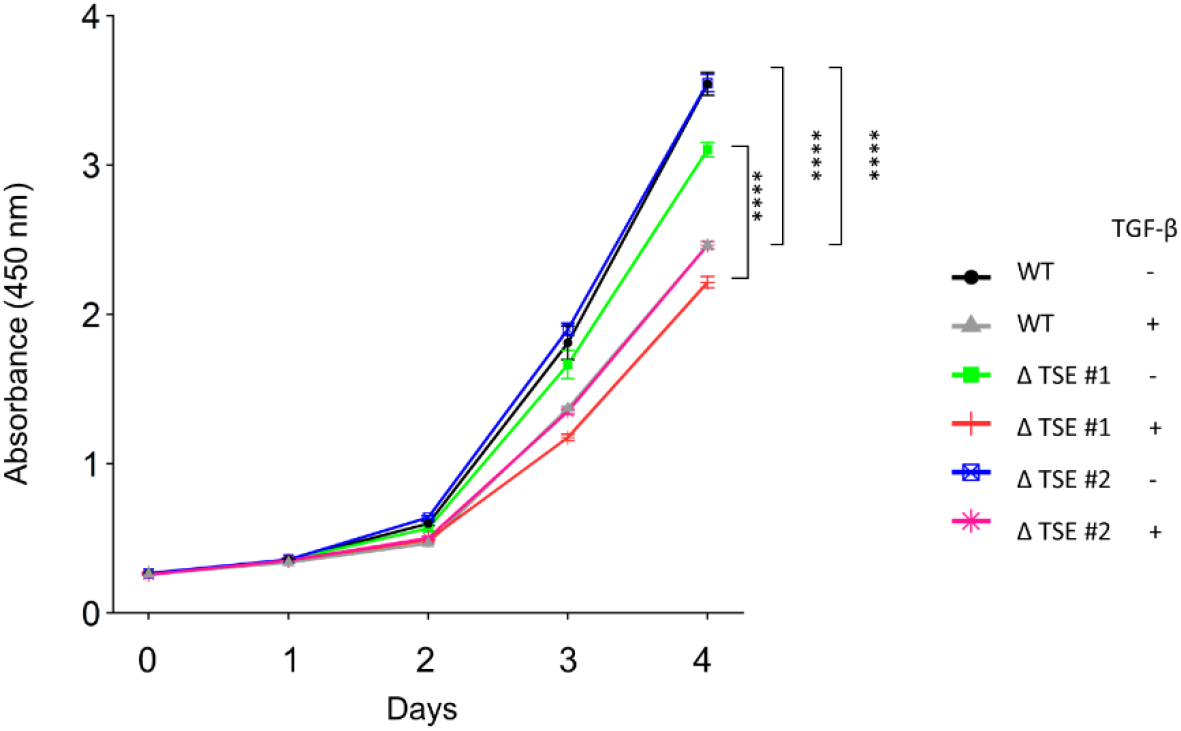
TGF-β reduced cell proliferation rate in both WT and ΔTSE cell lines. wild-type MCF10A cells and ΔTSE clones were treated with TGF-β1 (10 ng/ml). WST-8 was added at the indicated time points, and cells were incubated for 2h. Mean ratios ± SEs were obtained from five biologically independent samples. Representative result from three independent experiments are shown. ****, *p* < 0.0001; Unpaired two-tailed *t*-test.

**Figure S6.**
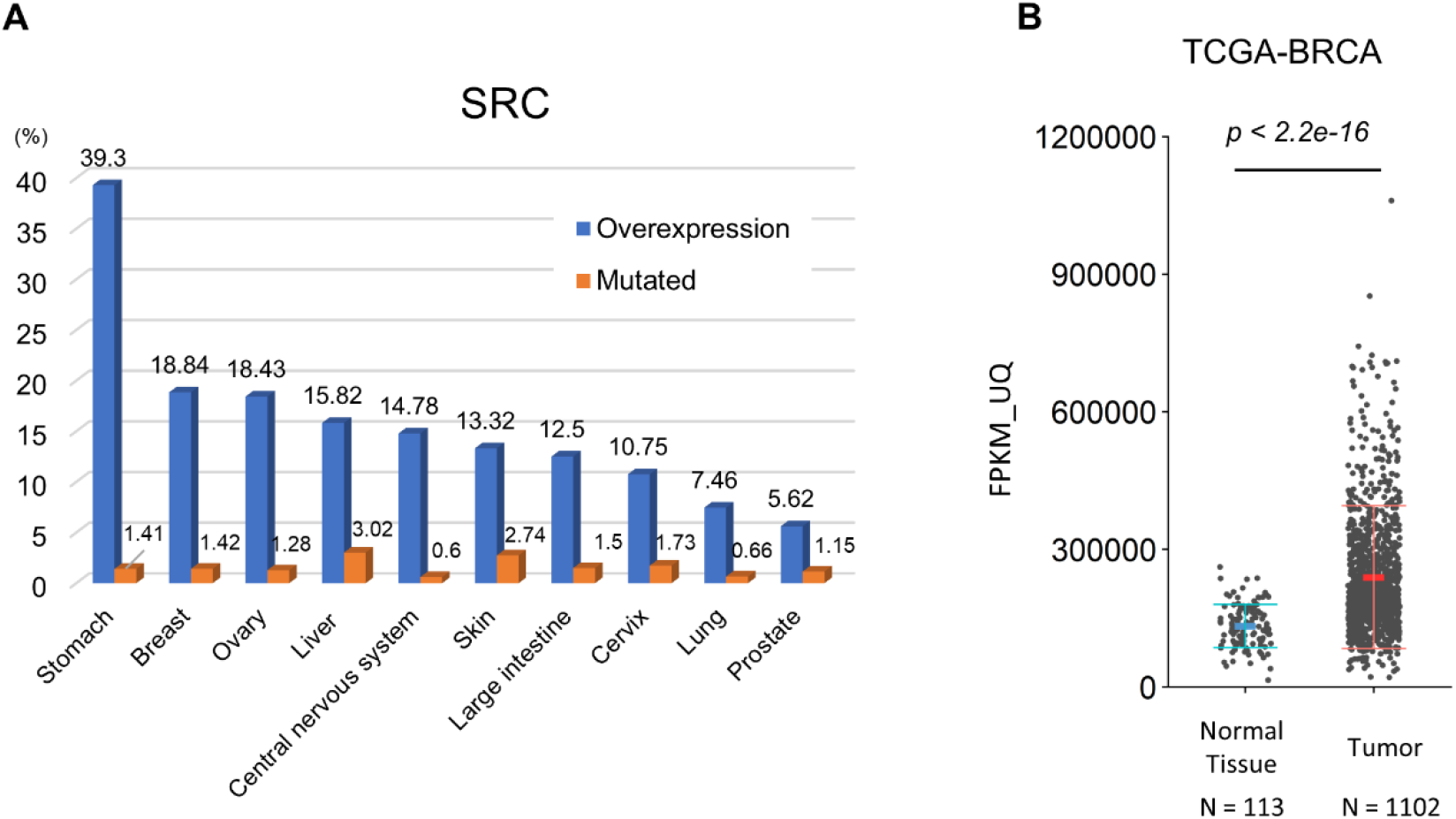
SRC is overexpressed in various types of human cancers. (A) The frequencies of somatic mutations in *SRC* in various human cancers were obtained from the Catalogue of Somatic Mutations in Cancer (COSMIC; https://cancer.sanger.ac.uk/cosmic). Blue and orange bars indicate the ratios of overexpression and genetic mutations, respectively. (B) SRC mRNA expression was analyzed using a cohort of breast cancer patients from The Cancer Genome Atlas (TCGA-BRCA). The *p*-value was determined using the Wilcoxon rank-sum test.

**Figure S7.**
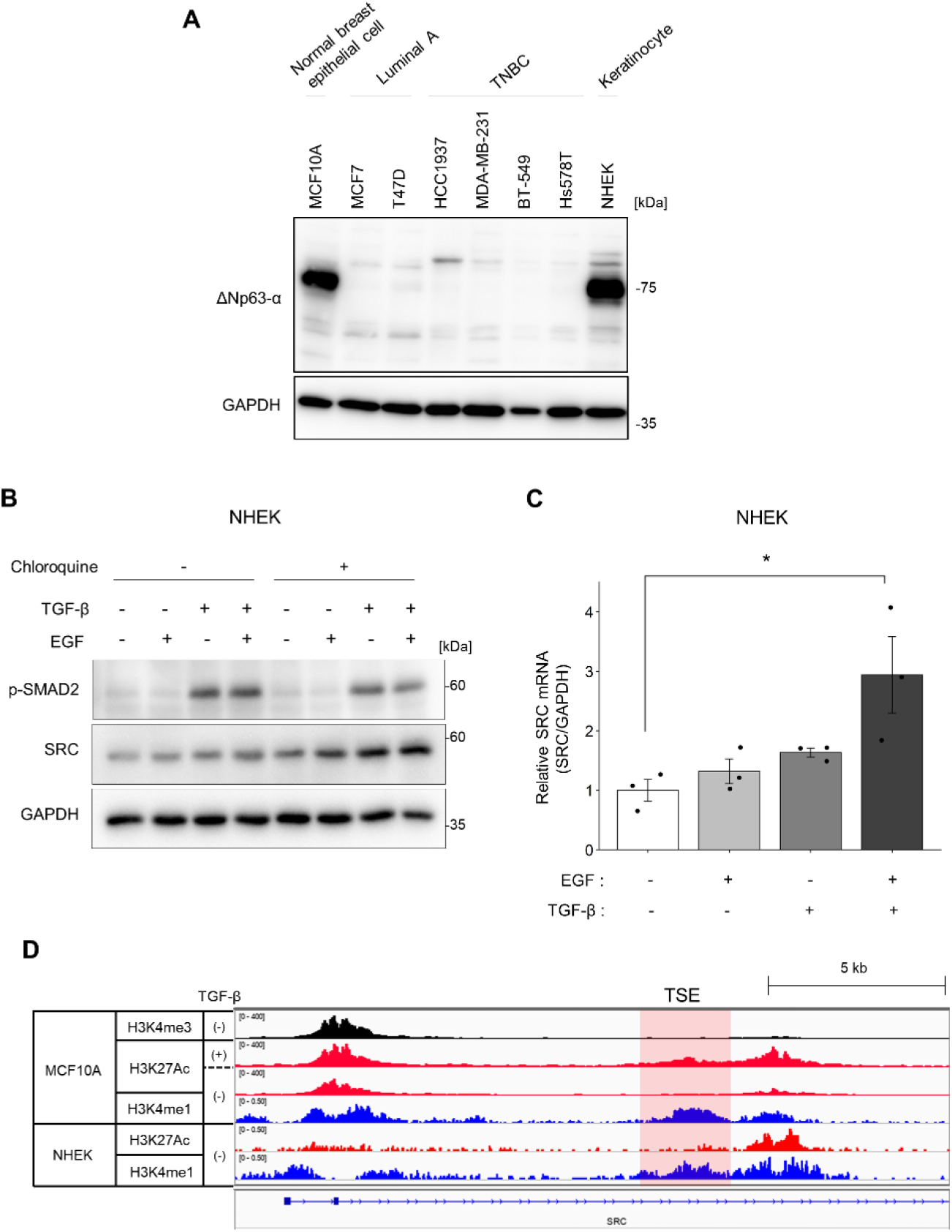
Co-stimulation of TGF-β and EGF regulates SRC expression in p63-expressing normal human epidermal keratinocyte (NHEK). (A) Immunoblot analysis for the expression of p63 in the indicated cell lines. TNBC, triple-negative breast cancer. (B) NHEK cells were treated with TGF-β1 (10 ng/ml) and/or EGF (20 ng/ml) for 24h. Chloroquine (100 μM) was added to the medium, and the cells were incubated for 2h before cell lysate collection. Cell lysates were subjected to immunoblotting using the indicated antibodies. (C) NHEK cells were treated with TGF-β1 (10 ng/ml) and/or EGF (20 ng/ml) for 24h. Total RNA was isolated and subjected to quantitative real-time polymerase chain reaction PCR. (D) Genomic loci of *the SRC* promoter region in MCF10A and NHEK cells The IP targets are H3K4me3 (black), H3K27Ac (red), and H3K4me1 (blue). (C) Mean ratios ± SEs were obtained from three independent experiments. * *p* < 0.05; One-way ANOVA with Tukey’s post hoc test.

**Supplementary Table S1.**
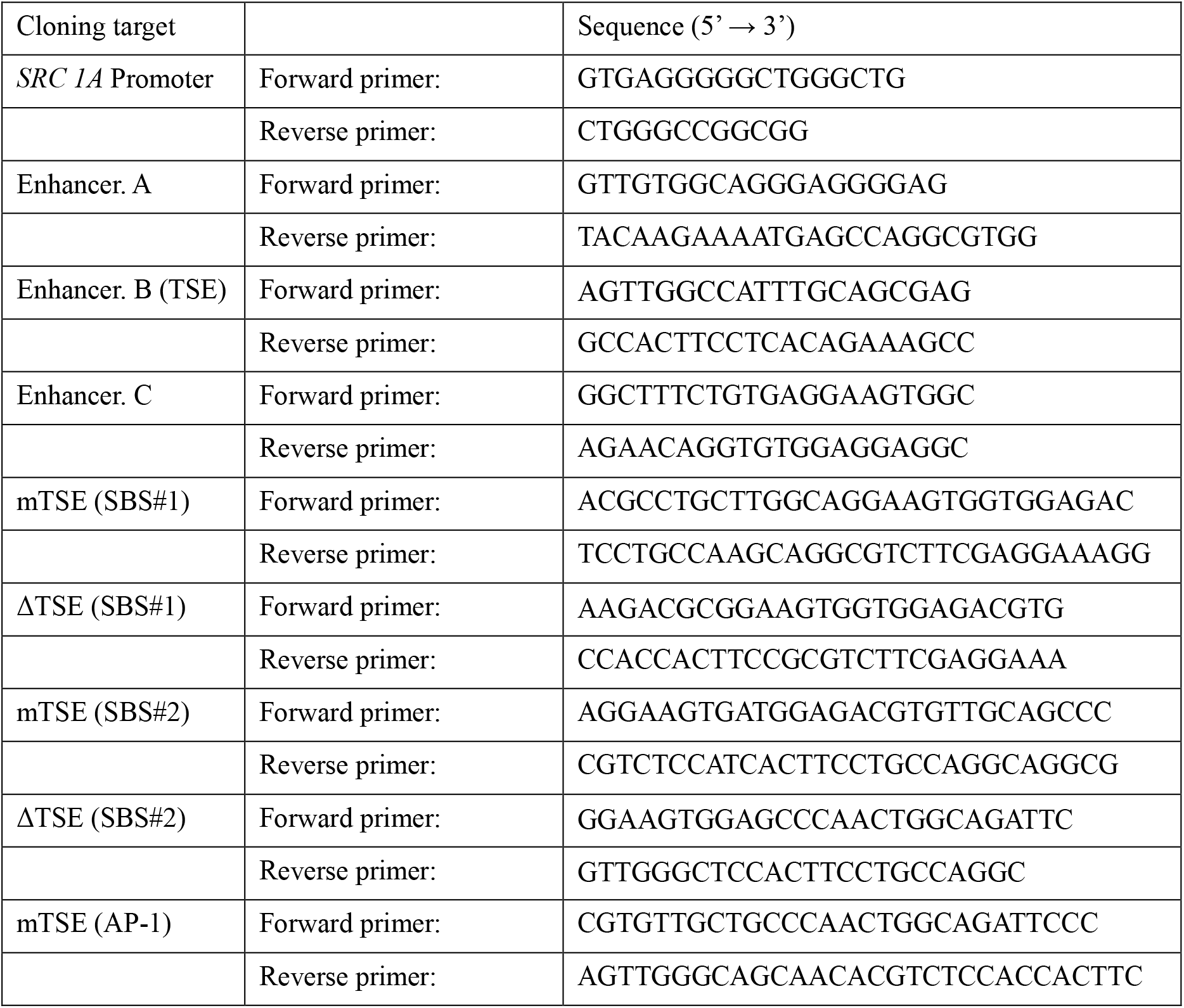
List of primers for luciferase reporter vector constructions.

**Supplementary Table S2.**
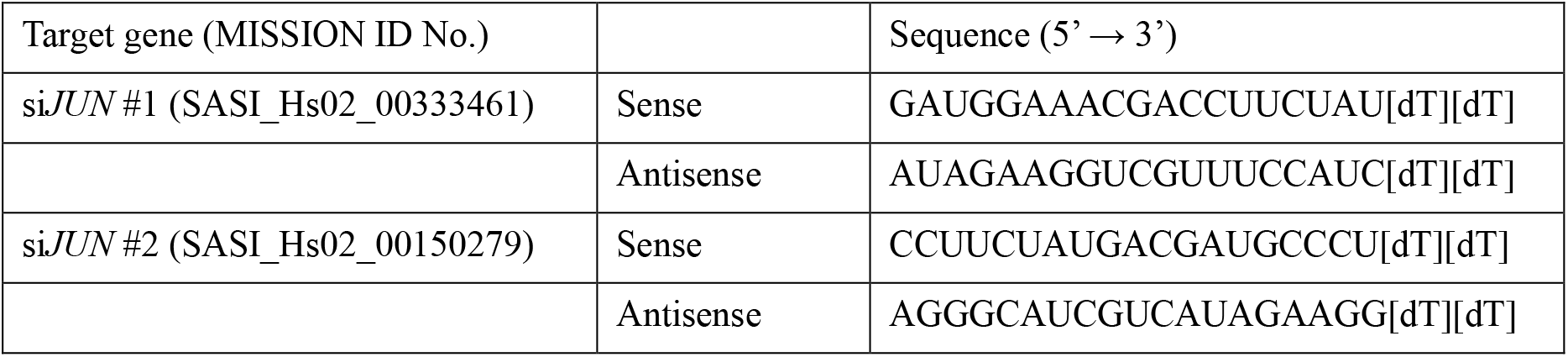
List of siRNA.

**Supplementary Table S3.**
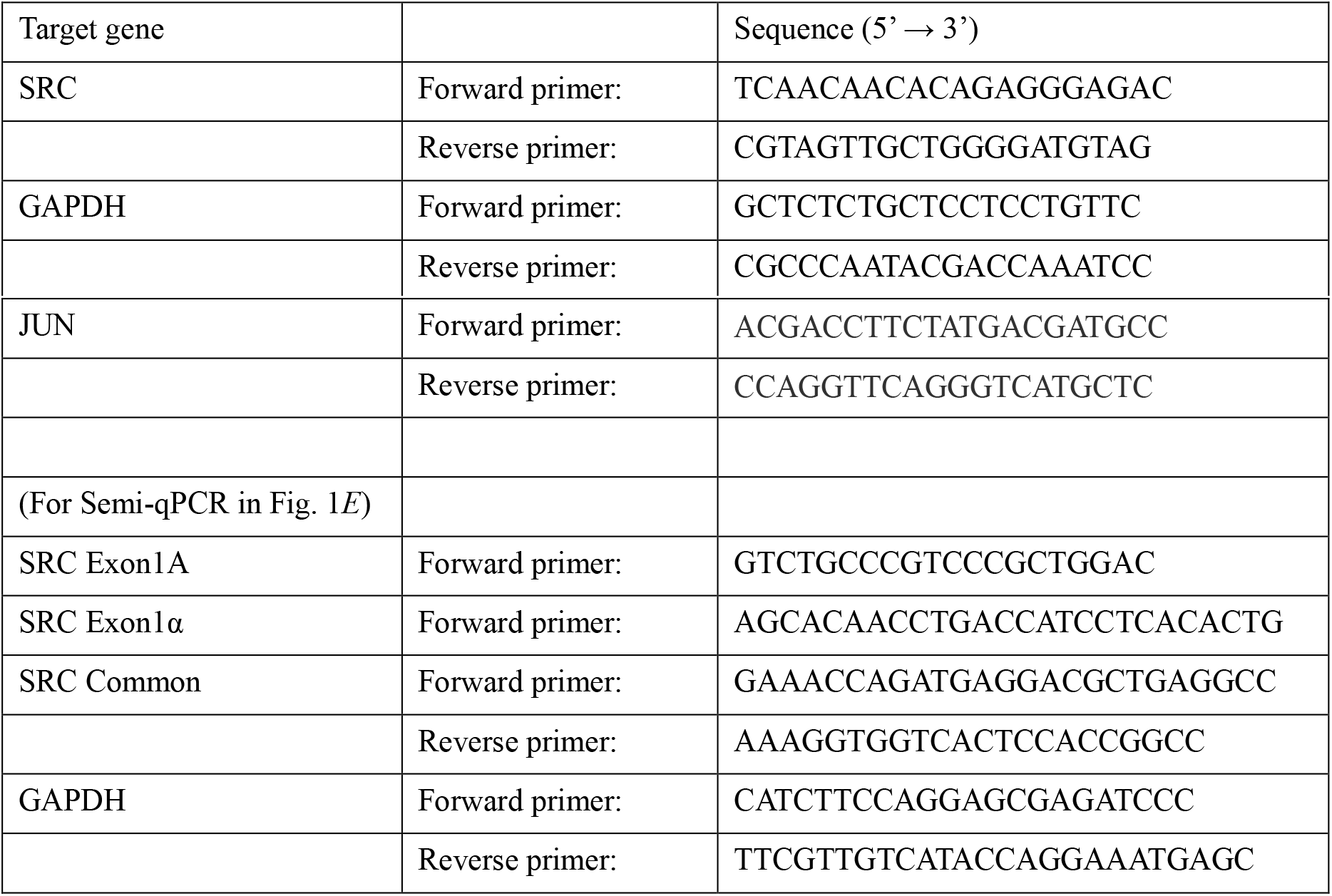
List of qPCR primers.

**Supplementary Table S4.**
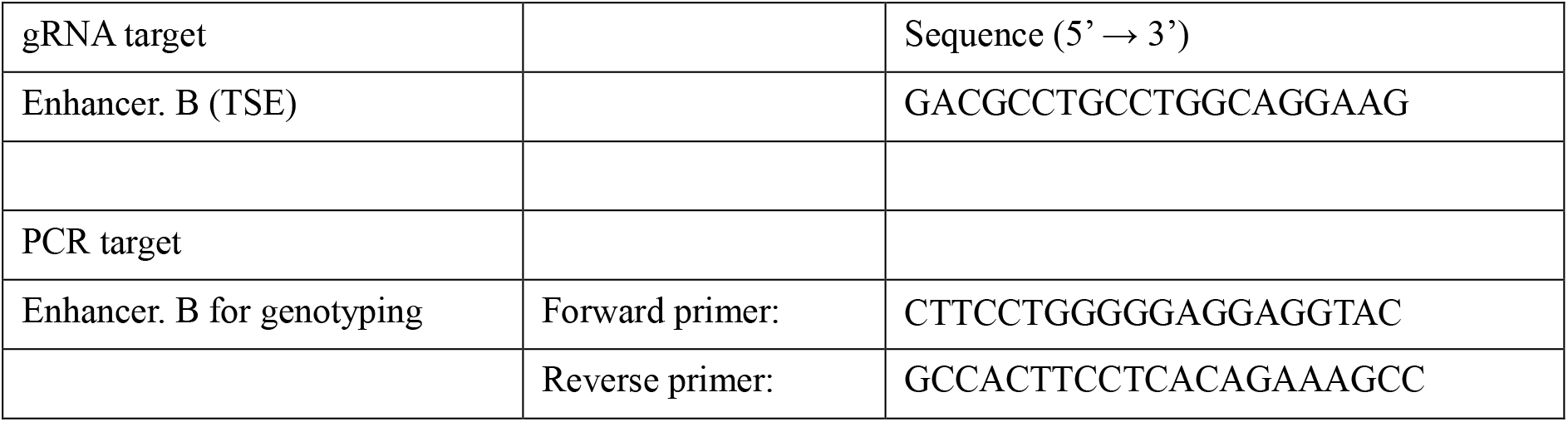
List of gRNA sequence and genotyping primers.

## Notes

### Competing Interest Statement

The authors have declared no competing interest.

### Summary of Updates

Protein markers has been added to all Western blotting data. Some of data and schematics has been rearranged to be more appropriate.

## References

Avizienyte, E. and Frame, M. C. (2005). Src and FAK signalling controls adhesion fate and the epithelial-to-mesenchymal transition. Curr. Opin. Cell Biol. 17, 542–547.

Avizienyte, E., Fincham, V. J., Brunton, V. G. and Frame, M. C. (2004). Src SH3/2 Domain-mediated Peripheral Accumulation of Src and Phospho-myosin Is Linked to Deregulation of E-cadherin and the Epithelial-Mesenchymal Transition. Mol. Biol. Cell 15, 2794–2803.

Bonham, K., Ritchie, S. A., Dehm, S. M., Snyder, K. and Boyd, F. M. (2000). Alternative, human SRC promoter and its regulation by hepatic nuclear factor-1α. J. Biol. Chem. 275, 37604–37611.

Castro-Mondragon, J. A., Riudavets-Puig, R., Rauluseviciute, I., Berhanu Lemma, R., Turchi, L., Blanc-Mathieu, R., Lucas, J., Boddie, P., Khan, A., Perez, N. M., et al. (2022). JASPAR 2022: The 9th release of the open-access database of transcription factor binding profiles. Nucleic Acids Res. 50, D165–D173.

Colak, S. and ten Dijke, P. (2017). Targeting TGF-β Signaling in Cancer. Trends in Cancer 3, 56–71.

Deckers, M., Van Dinther, M., Buijs, J., Que, I., Löwik, C., Van Der Pluijm, G. and Ten Dijke, P. (2006). The tumor suppressor Smad4 is required for transforming growth factor β-induced epithelial to mesenchymal transition and bone metastasis of breast cancer cells. Cancer Res. 66, 2202–2209.

Fujita, Y., Krause, G., Scheffner, M., Zechner, D., Leddy, H. E. M., Behrens, J., Sommer, T. and Birchmeier, W. (2002). Hakai, a c-Cbl-like protein, ubiquitinates and induces endocytosis of the E-cadherin complex. Nat. Cell Biol. 4, 222–231.

Fuxe, J., Mayor, R., Nieto, M. A., Puisieux, A., Runyan, R., Savagner, P., Thiery, J. P., Thompson, E. W., Theveneau, E. and Williams, E. D. (2020). Guidelines and definitions for research on epithelial–mesenchymal transition. Nat. Rev. Mol. Cell Biol.

Gaarenstroom, T. and Hill, C. S. (2014). TGF-β signaling to chromatin: How Smads regulate transcription during self-renewal and differentiation. Semin. Cell Dev. Biol. 32, 107–118.

Golubovskaya, V. M., Nyberg, C., Zheng, M., Kweh, F., Magis, A., Ostrov, D. and Cance, W. G. (2008). Site of Focal Adhesion Kinase Decreases Tumor Growth. 7405–7416.

Guarino, M. (2010). Src signaling in cancer invasion. J. Cell. Physiol. 223, 14–26.

Higuchi, M., Ishiyama, K., Maruoka, M., Kanamori, R., Takaori-Kondo, A. and Watanabe, N. (2021). Paradoxical activation of c-Src as a drug-resistant mechanism. Cell Rep. 34, 108876.

Ishizawar, R. and Parsons, S. J. (2004). C-Src and cooperating partners in human cancer. Cancer Cell 6, 209–214.

Joan Massagué (2008). TGFβ in Cancer. Cell 134, 215–230.

Kim, M., Tezuka, T., Tanaka, K. and Yamamoto, T. (2004). Cbl-c suppresses v-Src-induced transformation through ubiquitin-dependent protein degradation. Oncogene 23, 1645–1655.

Komiya, Y., Onodera, Y., Kuroiwa, M., Nomimura, S., Kubo, Y., Nam, J. M., Kajiwara, K., Nada, S., Oneyama, C., Sabe, H., et al. (2016). The Rho guanine nucleotide exchange factor ARHGEF5 promotes tumor malignancy via epithelial-mesenchymal transition. Oncogenesis 5,.

Kuroiwa, M., Oneyama, C., Nada, S. and Okada, M. (2011). The guanine nucleotide exchange factor Arhgef5 plays crucial roles in Src-induced podosome formation. J. Cell Sci. 1726–1738.

Le Coq, J., Acebrón, I., Rodrigo Martin, B., López Navajas, P. and Lietha, D. (2022). New insights into FAK structure and function in focal adhesions. J. Cell Sci. 135,.

Levy, L. and Hill, C. S. (2006). Alterations in components of the TGF-β superfamily signaling pathways in human cancer. Cytokine Growth Factor Rev. 17, 41–58.

Liberati, N. T., Datto, M. B., Frederick, J. P., Shen, X., Wong, C., Rougier-Chapman, E. M. and Wang, X. F. (1999). Smads bind directly to the Jun family of AP-1 transcription factors. Proc. Natl. Acad. Sci. U. S. A. 96, 4844–4849.

Liu, W., Guo, T. F., Jing, Z. T., Yang, Z., Liu, L., Yang, Y. P., Lin, X. and Tong, Q. Y. (2018). Hepatitis B virus core protein promotes hepatocarcinogenesis by enhancing Src expression and activating the Src/PI3K/Akt pathway. FASEB J. 32, 3033–3046.

López-Cortés, A., Paz-y-Miño, C., Guerrero, S., Cabrera-Andrade, A., Barigye, S. J., Munteanu, C. R., González-Díaz, H., Pazos, A., Pérez-Castillo, Y. and Tejera, E. (2020). OncoOmics approaches to reveal essential genes in breast cancer: a panoramic view from pathogenesis to precision medicine. Sci. Rep. 10, 1–21.

Maeda, M., Shintani, Y., Wheelock, M. J. and Johnson, K. R. (2006). Src activation is not necessary for transforming growth factor (TGF)-β-mediated epithelial to mesenchymal transitions (EMT) in mammary epithelial cells: PP1 directly inhibits TGF-β receptors I and II. J. Biol. Chem. 281, 59–68.

Majid, S., Saini, S., Dar, A. A., Hirata, H., Shahryari, V., Tanaka, Y., Yamamura, S., Ueno, K., Zaman, M. S., Singh, K., et al. (2011). MicroRNA-205 inhibits Src-mediated oncogenic pathways in renal cancer. Cancer Res. 71, 2611–2621.

Massagué, J. (2012). TGFβ signalling in context. Nat. Rev. Mol. Cell Biol. 13, 616–630.

Mayer, E. L. and Krop, I. E. (2010). Advances in targeting Src in the treatment of breast cancer and other solid malignancies. Clin. Cancer Res. 16, 3526–3532.

Mukhopadhyay, C., Triplett, A., Bargar, T., Heckman, C., Wagner, K. U. and Naramura, M. (2016). Casitas B-cell lymphoma (Cbl) proteins protect mammary epithelial cells from proteotoxicity of active c-Src accumulation. Proc. Natl. Acad. Sci. U. S. A. 113, E8228–E8237.

Nada, S., Okada, M., MacAuley, A., Cooper, J. A. and Nakagawa, H. (1991). Cloning of a complementary DNA for a protein-tyrosine kinase that specifically phosphorylates a negative regulatory site of p60c-src. Nature 351, 69–72.

Okada, M. (2012). Regulation of the Src family kinases by Csk. Int. J. Biol. Sci. 8, 1385–1397.

Okuzaki, D., Yamauchi, T., Mitani, F., Miyata, M., Ninomiya, Y., Watanabe, R., Akamatsu, H. and Oneyama, C. (2020). c-Src promotes tumor progression through downregulation of microRNA-129-1-3p. Cancer Sci. 111, 418–428.

Oneyama, C. and Okada, M. (2015). MicroRNAs as the fine-tuners of Src oncogenic signalling. J. Biochem. 157, 431–438.

Padua, D. and Massagué, J. (2009). Roles of TGFβ in metastasis. Cell Res. 19, 89–102.

Rb, I. and TJ, Y. (2000). Role of Src expression and activation in human cancer. Oncogene 19, 5636–5642.

Ritchie, S., Boyd, F. M., Wong, J. and Bonham, K. (2000). Transcription of the human c-Src promoter is dependent on Sp1, a novel pyrimidine binding factor SPy, and can be inhibited by triplex-forming oligonucleotides. J. Biol. Chem. 275, 847–854.

Shi, Y. and Massagué, J. (2003). Mechanisms of TGF-β signaling from cell membrane to the nucleus. Cell 113, 685–700.

Sundqvist, A., Morikawa, M., Ren, J., Vasilaki, E., Kawasaki, N., Kobayashi, M., Koinuma, D., Aburatani, H., Miyazono, K., Heldin, C. H., et al. (2018). JUNB governs a feed-forward network of TGFβ signaling that aggravates breast cancer invasion. Nucleic Acids Res. 46, 1180–1195.

Sundqvist, A., Vasilaki, E., Voytyuk, O., Bai, Y., Morikawa, M., Moustakas, A., Miyazono, K., Heldin, C. H., ten Dijke, P. and van Dam, H. (2020). TGFβ and EGF signaling orchestrates the AP-1-and p63 transcriptional regulation of breast cancer invasiveness. Oncogene 39, 4436–4449.

Tanaka, K., Ito, Y., Kajiwara, K., Nada, S. and Okada, M. (2020). Ubiquitination of Src promotes its secretion via small extracellular vesicles. Biochem. Biophys. Res. Commun. 525, 184–191.

Tsai, C. C., Kuo, F. T., Lee, S. B., Chang, Y. T. and Fu, H. W. (2019). Endocytosis-dependent lysosomal degradation of Src induced by protease-activated receptor 1. FEBS Lett. 593, 504–517.

Tsukamoto, T., Kajiwara, K., Nada, S. and Okada, M. (2019). Src mediates TGF-β-induced intraocular pressure elevation in glaucoma. J. Cell. Physiol. 234, 1730–1744.

Verbeek, B. S., Vroom, T. M., Adriaansen-Slot, S. S., Ottenhoff-Kalff, A. E., Geertzema, J. G. N., Hennipman, A. and Rijksen, G. (1996). c-Src protein expression is increased in human breast cancer. An immunohistochemical and biochemical analysis. J. Pathol. 180, 383–388.

Wang, N., Liang, H., Zhou, Y., Wang, C., Zhang, S., Pan, Y., Wang, Y., Yan, X., Zhang, J., Zhang, C. Y., et al. (2014). miR-203 suppresses the proliferation and migration and promotes the apoptosis of lung cancer cells by targeting SRC. PLoS One 9,.

Webb, D. J., Donais, K., Whitmore, L. A., Thomas, S. M., Turner, C. E., Parsons, J. T. and Horwitz, A. F. (2004). FAK-Src signalling through paxillin, ERK and MLCK regulates adhesion disassembly. Nat. Cell Biol. 6, 154–161.

Westhoff, M. A., Serrels, B., Fincham, V. J., Frame, M. C. and Carragher, N. O. (2004). Src-Mediated Phosphorylation of Focal Adhesion Kinase Couples Actin and Adhesion Dynamics to Survival Signaling. Mol. Cell. Biol. 24, 8113–8133.

Wiener, J. R., Windham, C., Estrella, V. C., Parikh, N. U., Thall, P. F., Deavers, M. T., Bast, R. C., Mills, G. B. and Gallick, G. E. (2003). Activated Src protein tyrosine kinase is overexpressed in late-stage human ovarian cancers. Gynecol. Oncol. 88, 73–79.

Yeatman, T. J. (2004). A renaissance for SRC. Nat. Rev. Cancer 4, 470–480.

Zientek-Targosz, H., Kunnev, D., Hawthorn, L., Venkov, M., Matsui, S. I., Cheney, R. T. and Ionov, Y. (2008). Transformation of MCF-10A cells by random mutagenesis with frameshift mutagen ICR191: A model for identifying candidate breast-tumor suppressors. Mol. Cancer 7, 1–12.

